# Loss of endothelial ALK1 signaling induces the emergence of a KIT+ angiogenic endothelial cluster driving brain arteriovenous malformations

**DOI:** 10.1101/2025.06.05.657957

**Authors:** Elise Drapé, Lauranne Carrier, Gael Cagnone, Halima Drissi Touzani Walali, Mathilde Bizou, Patrick Piet Van Vliet, Gregor Andelfinger, Bruno Larrivée, Alexandre Dubrac

## Abstract

**Background:** Hereditary Hemorrhagic Telangiectasia type 2 (HHT2) is a genetic disorder caused by mutations in the *ALK1* (*ACVRL1*) gene, encoding a receptor for Bone Morphogenetic Proteins 9 and 10 (BMP9/BMP10). HHT2 patients frequently develop brain arteriovenous malformations (bAVMs), which are abnormal connections between arteries and veins. Currently, surgical resection is the only treatment, associated with significant risks and complications. Despite evidence suggesting endothelial cell (EC) heterogeneity in bAVMs, it remains poorly characterized, limiting our ability to identify new therapeutic avenues.

**Methods:** We employed endothelial cell-specific and inducible *Alk1* knockout mice (*Alk1iECKO*) with tamoxifen-induced deletion at postnatal day 6 (P6). We separately analyzed the P8 perineural (PNVP) and intraneural (INVP) vascular plexuses, which differ in vessel composition and flow dynamics. Single-cell RNA sequencing (scRNAseq) was performed to characterize EC heterogeneity and identify transcriptomic changes in both vascular plexuses of mutant versus wild type mice.

**Results:** Loss of endothelial ALK1 signaling triggered bAVM formation predominantly in the PNVP vascular network. scRNAseq revealed that *Alk1* deletion promoted brain capillaries differentiation into angiogenic-1 ECs, whereas it drove PNVP venules EC proliferation and the emergence of the unique angiogenic-2 cluster. The latter shares transcriptomic features with human AVM ECs, including angiogenic tip cell markers and a strong glycolytic signature. Among its defining markers, *Kit* emerged as a direct downstream target of BMP9-ALK1 signaling. Pharmacological KIT inhibition using Masitinib, Imatinib, or KIT-blocking antibodies prevented bAVM formation in *Alk1iECKO* mice.

**Conclusion:** Our study uncovers a previously unrecognized EC population, the angiogenic-2 cluster, as a key contributor to bAVM development. We identify *Kit* as a central regulator of this cluster, establishing it as a promising therapeutic target for preventing bAVMs in HHT2.

**Clinical Perspective:** *What is new?:*

- Using endothelial-specific *Alk1* knockout mouse models and single-cell transcriptomics, we identified a novel angiogenic endothelial cell population as a key driver of brain AVM formation.
- This angiogenic EC cluster shares molecular features with human bAVM cells, including high expression of *KIT*, which we identified as a new direct transcriptional target of BMP9-ALK1 signaling.
- Pharmacological inhibition of KIT using small molecules or blocking antibodies effectively prevents AVM formation *in vivo*, establishing KIT as a promising therapeutic target.

*What are the clinical implications?:*

- Our findings support the therapeutic potential of targeting KIT with FDA-approved drugs such as Imatinib to treat HHT2-associated brain AVMs, offering a non-invasive alternative to surgical intervention.
- Characterizing AVM-specific endothelial subtypes may enable the development of targeted and personalized therapies, improving patient outcomes and minimizing treatment-associated risks in HHT.

---

Hereditary Hemorrhagic Telangiectasia (HHT), or Rendu-Osler syndrome, is a rare genetic disorder affecting 1 in 5,000 to 8,000 individuals worldwide^1,2^. It is caused by mutations in components of the BMP9/ALK1 signaling pathway, which regulates vascular development and homeostasis. Mutations in the *ENG* and *ACVRL1* genes, encoding ENDOGLIN and ALK1, respectively, account for the majority of HHT cases (HHT1 and HHT2) ^3–5^. Rare mutations in *GDF2* or *SMAD4* result in HHT5 and juvenile polyposis-HHT (JP-HHT)^6,7^. HHT patients develop recurrent epistaxis, telangiectasia, and arteriovenous malformations (AVMs) in various organs, with brain AVMs (bAVMs) occurring in 10–20% of cases ^8,9^. AVMs are vascular anomalies characterized by the direct connection of arteries and veins without an intervening capillary bed. These lesions are associated with severe complications, including epilepsy and cerebral hemorrhage^10,11^. Current management relies on high-risk surgical interventions, and pharmacological treatments for bAVMs are limited, underscoring the need for new therapeutic approaches^12,13^.

The pathogenesis of bAVM remains poorly understood. BMP9 and BMP10 bind ALK1 and its co-receptors, including BMPRII and ENDOGLIN, activating SMAD1/5/8 and SMAD4 transcription factors. Studies using different HHT mouse models have highlighted that deletions of ALK1, ENDOGLIN, SMAD1/5, or SMAD4 in ECs are sufficient to induce bAVMs^14–17^. Moreover, recent studies have demonstrated that the deletion of *ALK1* and *SMAD4,* specifically in venous ECs induces AVM formations, whereas similar deletions in arterial endothelial cells do not, suggesting that AVMs originate primarily from venous endothelial cells^18,19^. Loss of BMP9/ALK1/SMAD4 signaling promotes shear stress, activating VEGF-induced PI3K-AKT signaling and the integrin/YAP/TAZ pathways^16,18–21^. As a result, mutant ECs display hyperproliferation, loss of arteriovenous identity, and impaired migration against blood flow ^16,18,19,22,23^. However, most mechanistic insights have been derived from postnatal retinal models, while the specific cellular and molecular mechanisms in bAVMs remain poorly understood. Moreover, these findings suggest the BMP9/ALK1/SMAD4 signaling disruptions alter the transcriptomic landscape of ECs, driving cellular heterogeneity and contributing to the diverse phenotypes observed in AVMs.

In addition to these molecular mechanisms, EC heterogeneity has emerged as a defining feature of AVMs, including those in the brain. This heterogeneity encompasses differences in cellular identity, metabolic activity, and proliferative states, all of which contribute to the distinct structural and functional abnormalities of AVMs. Pathological endothelial plasticity, characterized by the presence of proliferative ECs with mixed arteriovenous characteristics, disrupts physiological arteriovenous zonation. Recent studies utilizing single-cell transcriptomics have provided critical insights into this heterogeneity. For instance Winkler and colleagues^24^ identified a distinct subset of ECs in brain AVMs exhibiting heightened angiogenic and immunogenic potential and originating from the AVM nidus. Moreover, this pathological remodeling involves alterations in CNS-specific vascular properties coupled with the reactivation of developmental pathways^25^. Similarly, a recent study in retinal *Alk1* mutant mice has shown that some ECs within AVMs exhibit enhanced proliferation and mixed arteriovenous characteristics^26^. Yang et *al*^27^ further described arterial-lymphatic-like ECs arising from arterial ECs after *Alk1* deletion, implicating aberrant plasticity in lung AVM development. However, the specific EC populations driving HHT AVM pathogenesis in the brain and retina remain undefined. Identifying and characterizing EC driving AVM is essential for understanding AVM pathogenesis and developing effective therapeutic strategies.

Here, we investigate the contribution of EC heterogeneity to brain AVM pathogenesis in HHT2 using endothelial-specific *Alk1* knockout mouse model and single-cell transcriptomics. We identify distinct angiogenic EC populations, including a previously uncharacterized cluster that emerges specifically in the brain surface vasculature and exhibits transcriptomic similarity to human bAVM ECs. Among the defining markers of this population, we uncover KIT as a direct downstream target of BMP9-ALK1 signaling. Moreover, pharmacological KIT signaling inhibition effectively prevents AVM formation in vivo. Our findings highlight a unique angiogenic EC population as a key driver of AVM formation in HHT2 and establish KIT as a promising therapeutic target for the treatment of brain AVMs.

## Methods

### Animals models and experiments

*Alk1* deletion in endothelial cells (ECs) was achieved by *Alk1 l/l* mice with *Cdh5CREERT2* mice to generate *Alk1iECKO* mice. Postnatal deletion was induced via intraperitoneal injection of tamoxifen (Sigma, T5648; 15 µg) at postnatal day 6 (P6). Cre-negative littermates *Alk1 l/l* treated with tamoxifen were used as controls. Imatinib Mesylate (100 mg/kg, Selleckchem, S1026), Masitinib Mesylate (100mg/Kg, Cerdalane, E0814), KIT blocking antibody (500 µg, *InVivo*MAb anti-mouse c-Kit, BioxCell, BE0293-5MG-A) were administered intraperitoneally at P6, 3 hours following tamoxifen injection, and again at P7. For the *in vivo* proliferation assay, mice received an intraperitoneal injection of EdU (30 mg/kg; Life Technologies Inc., A10044) 3 hours prior to sacrifice. Mice were euthanized at P8. All breeding colonies and experimental protocols were approved by the Institutional Committee on Animal Research Ethics (CIBPAR).

### Sample Fixation and Preparation

After euthanasia, brains were fixed in a 4% paraformaldehyde solution (Fisher Scientific, AAJ19943K2) at 4°C overnight.

#### PNVP Staining

The cerebral cortexes were separated, then directly permeabilized and blocked in a blocking buffer containing 0.5% Triton X-100 (Sigma, X100-500ML), 1% BSA (Bioshop, ALB001.100), and 4% donkey serum (Fisher Scientific, 56-646-05ML) at 4°C overnight. Primary antibodies (**Table 1**) were added directly to the blocking solution and incubated for 48 hours at 4°C. After three PBS washes, secondary antibodies (**Table 2**) were applied in the blocking solution (1:500 dilution) and incubated for 24 hours at 4°C. Following three additional washes, the cortixes were placed on an agarose mold (**Figure 1A**) and mounted with mounting media (Sigma, 81381-50G). Imaging was performed directly with BioTek Lionheart FX Automated Microscope.

**Figure 1:**
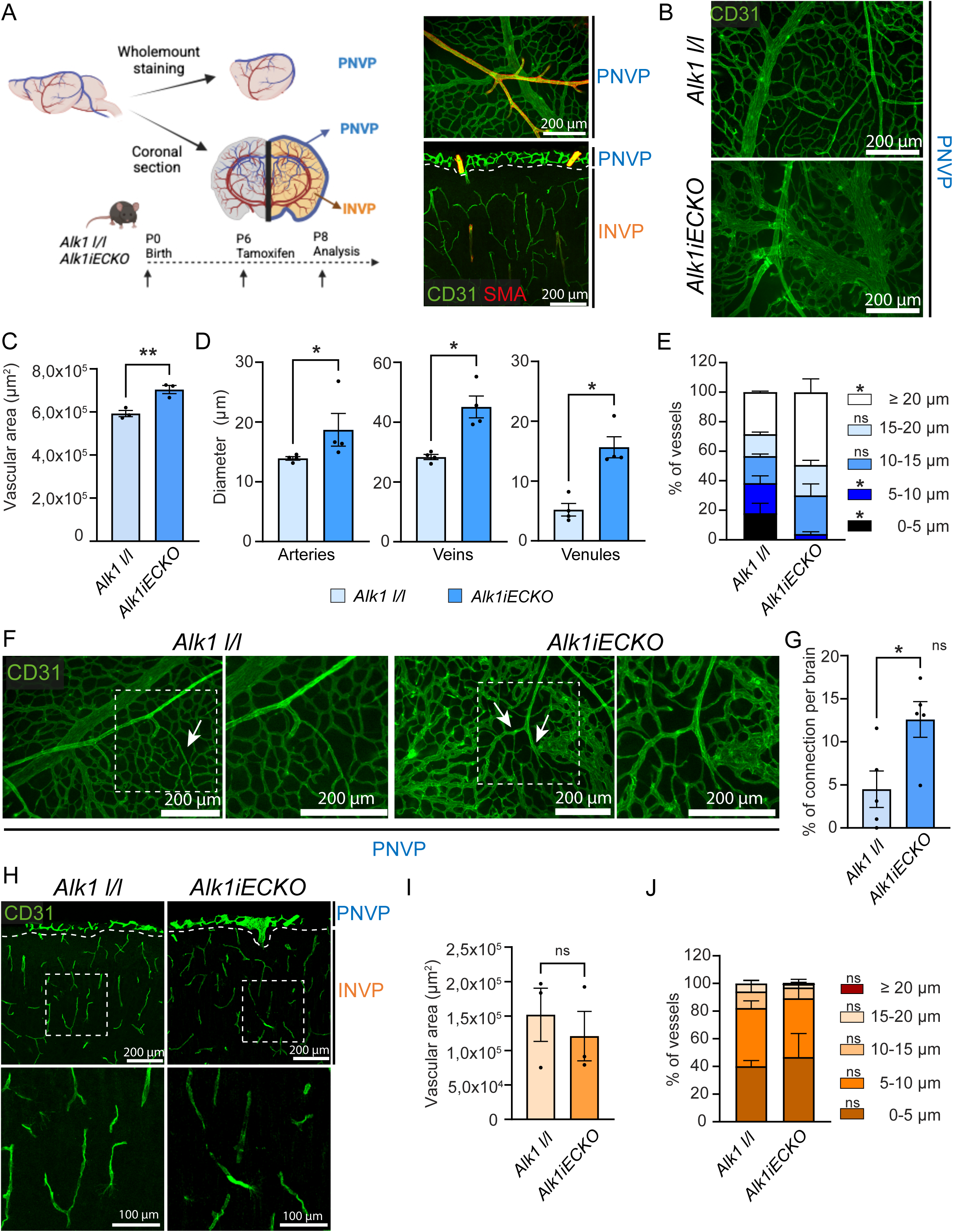
Loss of Alk1 signaling induces vascular malformations in the PNVP. **A**, Left, experimental timeline of tamoxifen administration and tissue analysis in *Alk1iECKO* and control mice. Right, representative images showing CD31 and aSMA immunostaining of the PNVP and INVP at P8, indicating vascular morphology. **B**, CD31 immunostaining of the PNVP of *Alk1iECKO* and control mice at P8. **C,** Quantification of vascular area in the PNVP of *Alk1iECKO* and control mice. **D-E,** Quantification of vessel diameter in the PNVP of *Alk1iECKO* and control mice at P8. **F**, CD31 immunostaining showing direct arteriovenous connections in the PNVP of *Alk1iECKO* mice. **G**, Quantification of PNVP AV shunts in the PNVP of *Alk1iECKO* and control mice. **H**, CD31 immunostaining of brain section from *Alk1iECKO* and control mice at P8. **I-J**, Quantification of vascular area (**I**) and vessel diameter (**J**) in the INVP of *Alk1iECKO* and control mice at P8. Each dot represents one mouse. Error bars represent means ± s.e.m, *\*P<0.05, **P<0.01,* Mann-Whitney test, Welch’s t test (C, D) or unpaired t test (E, F, I, J) were performed.

**Table 1.**
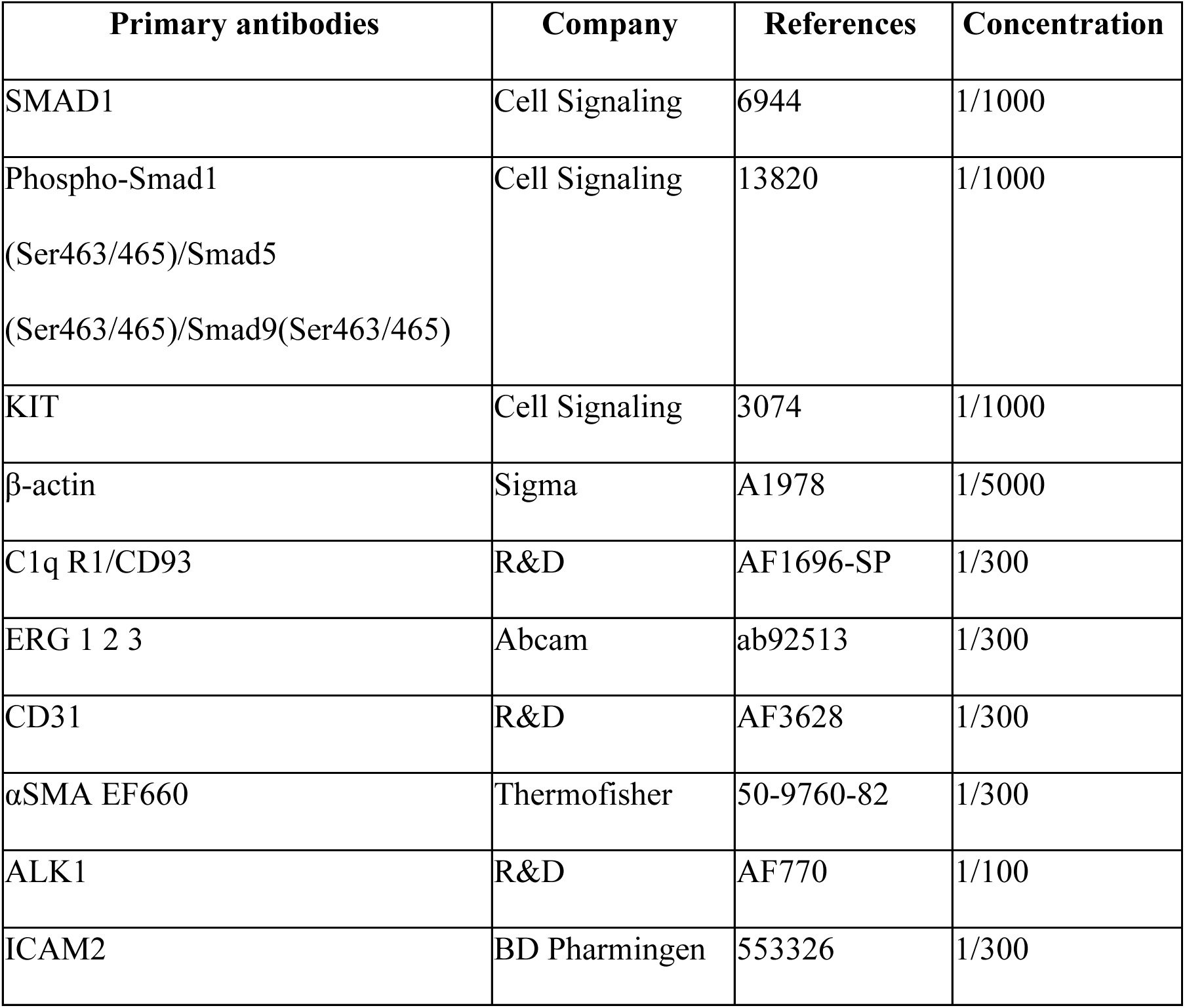

| Primary antibodies | Company | References | Concentration |
| --- | --- | --- | --- |
| SMAD1 | Cell Signaling | 6944 | 1/1000 |
| Phospho-Smad1<br>(Ser463/465)/Smad5<br>(Ser463/465)/Smad9(Ser463/465) | Cell Signaling | 13820 | 1/1000 |
| KIT | Cell Signaling | 3074 | 1/1000 |
| β-actin | Sigma | A1978 | 1/5000 |
| C1q R1/CD93 | R&D | AF1696-SP | 1/300 |
| ERG 1 2 3 | Abcam | ab92513 | 1/300 |
| CD31 | R&D | AF3628 | 1/300 |
| αSMA EF660 | Thermofisher | 50-9760-82 | 1/300 |
| ALK1 | R&D | AF770 | 1/100 |
| ICAM2 | BD Pharmingen | 553326 | 1/300 |

**Table 2.**

| Secondary antibodies | Company | References | Concentration |
| --- | --- | --- | --- |
| Anti-rabbit IgG, HRP-linked Antibody | Cell Signaling | 7074S | 1/10000 |
| Mouse anti-Goat IgG, HRP, Species<br>adsorbed, Polyclonal Secondary Antibody | Fisher | AP186PMI | 1/10000 |
| Donkey anti-Sheep IgG (H+L) Cross-Adsorbed Secondary Antibody, Alexa Fluor™ 488 | Thermofisher | A-11015 | 1/500 |
| Donkey anti-Rabbit IgG (H+L) Highly Cross-Adsorbed Secondary Antibody, Alexa Fluor™ 568 | Thermofisher | A10042 | 1/500 |
| Donkey anti-Goat IgG (H+L) Cross-Adsorbed Secondary Antibody, Alexa Fluor™ 488 | Thermofisher | A11055 | 1/500 |
| Chicken anti-Rat IgG (H+L) Cross-Adsorbed Secondary Antibody, Alexa Fluor™ 647 | Thermofisher | A21472 | 1/500 |

#### INVP Staining

Brains were immersed in a 30% sucrose (Fisher, BP220 1), solution for 48 hours, then embedded and frozen in OCT (Fisher Scientific, 23-730-571). The immunofluorescence protocol was conducted on 30 μm thick floating brain sections. Brain sections were permeabilized and blocked overnight in a blocking buffer containing 0.5% Triton X-100, 1% BSA, and 4% donkey serum. Primary antibodies (**Table 1**) were then incubated in the blocking buffer for 24 hours, followed by PBS washes and a 4-hour incubation with secondary antibodies (**Table 2**) in the blocking buffer at room temperature. After washing with PBS, brain slices were incubated for 5 minutes with DAPI at a concentration of 2 µg/mL (Sigma, D9542), then mounted using a fluorescent mounting media.

#### Proliferation assay

3 hours before mice sacrifice mice were intraperitoneally injected with 30mg/kg of EdU (Life Technologies Inc., A10044, stock solution at 5mg/mL). The click it EdU cell proliferation kit (Invitrogen, C10340) was performed on 100µm thick brain sections followed by ERG123 (Abcam, ab92513) immunostaining. Double positive nucleus staining was counted in order to determine the proportion of the proliferative endothelial cells.

#### In situ hybridization

Floating brain sections were washed 3 times in PBS, mounted on Superfrost Plus microscope slides (Fisher Scientific, 12-550-15), dry 10min at RT and baked 30min at 60°C. Sample were fixed 15min in 4% PFA and dehydrated with ethanol baths of gradually increasing concentration, dried, incubated with hydrogen peroxide during 10min and with protease IV during 30min. Probes hybridization were made during 2h at 40°C and for amplification slide were incubated with RNAscope Multiplex FL v2 Amp 1 and then Amp 2 for 30 min at 40°C and finally with RNAscope Multiplex FL v2 Amp 3 for 15 min at 40°C. Then, depending on each probe, slides were incubated with the appropriate RNAscope Multiplex FL v2 HRP-C1 or 2 or 3 for 15 min at 40C, washed twice with Wash Buffer 2 min at RT, then incubated with the appropriate fluorophore and incubated with RNAscope Multiplex FL v2 HRP blocker for 15 min at 40°C. After washing the slides for 2 min at RT, the slides were CD31 immunofluorescence stained and mounted as previously described. Brain slides were imaged with a sp8 confocal (Leica TCS SP8 laser scanning confocal microscope).

Floating brain sections were washed three times in PBS and mounted on Superfrost Plus microscope slides (Fisher Scientific, 12-550-15). The slides were air-dried for 10 minutes at room temperature (RT) and baked for 30 minutes at 60°C. Samples were fixed in 4% paraformaldehyde (PFA) for 15 minutes and dehydrated through a series of ethanol baths with gradually increasing concentrations. After drying, sections were treated with hydrogen peroxide for 10 minutes followed by incubation with Protease IV for 30 minutes. Probe hybridization was carried out at 40°C for 2 hours using RNAscope probes (**Table 3**). Signal amplification was performed by sequential incubation with RNAscope Multiplex FL v2 Amplification 1 and 2 reagents for 30 minutes each at 40°C, followed by RNAscope Multiplex FL v2 Amplification 3 for 15 minutes at 40°C. Depending on the specific probe, slides were then incubated with the appropriate RNAscope Multiplex FL v2 HRP-C1, HRP-C2, or HRP-C3 reagent for 15 minutes at 40°C. Slides were washed twice for 2 minutes at RT using Wash Buffer and incubated with the corresponding fluorophore before the incubation with RNAscope Multiplex FL v2 HRP Blocker for 15 minutes at 40°C. Following final washes, slides underwent immunofluorescence staining for CD31 and were mounted as previously described. Imaging was performed using a Leica TCS SP8 laser scanning confocal microscope.

**Table 3.**

| **RNASCOPE probes** | **Company** | **References** |
| --- | --- | --- |
| Mm Kit C2 | ACD | 314151 |
| Mm Stc1 C1 | ACD | 468371 |
| Mm Slc16a3 C3 | ACD | 432931 |

### Brain preparation for Single-cell RNA sequencing

Brain from mutant mice and littermate controls (*Alk1l/l* P8 N=2; *Alk1iECKO* P8 N=2) were collected and directly immersed and digested using the Neural Tissue Dissociation Kit (P) (Miltenyi Biotec, 130-092-628). Each brain was digested at 37°C for 30 minutes. During the enzymatic reaction, gentle flushing of the brain tissue was performed to separate the PNVP from the rest of the brain. The reaction was stopped by adding DMEM containing 5% fetal bovine serum (FBS), and the enzymatic solution containing the PNVP was filtered through a 40 μm cell strainer (Falcon, 352340). The remaining brain tissue was rinsed with PBS, mechanically dissociated, and subjected to a second round of enzymatic digestion. After further homogenization by pipetting, digestion was stopped with 5% FBS in DMEM, and the suspension was again passed through a 40 μm cell strainer. Myelin and cell debris were removed using the Debris Removal Solution (Miltenyi Biotec, 130-109-398), which separates components into distinct phases. Red blood cells were lysed with Red Blood Cell Lysis Solution (Miltenyi Biotec, 130-094-183). The resulting cell suspension was washed with a FACS buffer (PBS containing 1% FBS and 0.2% EDTA). Cells were stained for 30 minutes at room temperature with PE anti-mouse CD140b Antibody (Biolegend, 136006); APC-conjugated anti-mouse CD31 antibody (BD Biosciences, 551262) and FITC-conjugated anti-mouse CD45 antibody (BioLegend, 103108) in Fc block solution (5 µg/mL CD16/CD32 Fc Block, eBioscience, 16-0161-82). After washing, cells were stained with Live/Dead Fixable Aqua Dead Cell Stain (ThermoFisher Scientific, L34957) for 20 minutes at room temperature. Stained cells were washed and resuspended in PBS. The gating strategy was defined using unstained and single-stained controls. Dead cells and doublets were excluded, and endothelial cells (CD45⁻ CD31⁺) were sorted using a BD FACS Aria Fusion. Cells were resuspended in PBS containing 0.04% BSA and subjected to single-cell RNAseq.

#### Single-cell Transcriptomic analysis

Four libraries (*Alk1 l/l* PNVP, *Alk1 l/l* INVP, *Alk1iECKO* PNVP, *Alk1iECKO* INVP) were generated using the Chromium Single Cell 3’ Reagent Kits v3.1 (10x Genomics; Pleasanton, CA, USA). Libraries were sequenced on Illumina NovaSeq6000 S4 (mean of ∼144 00 read/cell), followed by de-multiplexing and mapping to the mouse genome (Mus_musculus:GRCm38) using CellRanger (cellranger-6.1.1, 10x Genomics). Count matrices were analyzed using Seurat V5^28^. Doublets were identified using the Solo package (https://www.cell.com/cell-systems/fulltext/S2405-4712(20)30195-2) and removed from further analysis. Genes expressed in less than 10 cells were excluded. We kept cells with more than 200 genes and less than 6500 genes, less than 45000 transcripts and less than 10% mitochondrial transcripts. Following quality control and initial dimensionality reduction, endothelial cells were then subclustered and analyzed separately. Briefly, UMAP-identified ECs within each of the 4 four datasets (*Alk1 l/l* PNVP = 1195 ECs, *Alk1 l/l* INVP = 3221 ECs, *Alk1ECKO* PNVP = 2193 ECs, *Alk1iECKO* INVP = 1904 ECs) were merged together (mean nFeature/cell = 4097, mean nCount/cell = 16763), normalized using SCTranform (regression on nFeature_RNA and percent.mt) then dimensionality reduction was performed using runUMAP based on the first 20 PCs from PCA analysis of the top 3000 most variable genes (SCT assay). Clustering was then performed on UMAP first and second dimensions at a resolution of 1.2, followed by differential marker expression analysis between clusters based on normalised RNA count (RNA assay). Normalised RNA counts were then scaled (regression on nFeature_RNA and percent.mt) for data visualization. Clusters were finally annotated based on known endothelial subtype marker genes. Gene expression from the RNA assay was visualised using featureplot or dotplot functions. Pathway activity score computation using the MsgiDB database (msigdb.v2023.1.Mm.symbols.gmt) or in house datasets was performed based on gene counts from the SCT assay using the package VISION (https://www.nature.com/articles/s41467-019-12235-0). Differential gene expression or pathway score was done using the Wilcoxon rank sum test and visualised on a log2 fold-change basis. Trajectory analysis was performed with Monocle3 package (https://cole-trapnell-lab.github.io/monocle3/) using normalised RNA counts and precomputed UMAP dimensions from Seurat analysis. Pseudotime analysis was performed on Monocle3 graph trajectory using the vein subtype as root. Cell-cell communication (CCC) analysis was performed using the Connectome package (https://doi.org/10.1038/s4022-07959-x) on the RNA assay, computing a z-score for each ligand-receptor pair between cell types. CCC was visualized using circos plot using thresholds on percent of ligand and receptor expression in each cell types as well as ligand-receptor z-score. Public dataset from Winkler et al and Wälchli^24,25^ et al was downloaded (https://adult-brain-vasc.cells.ucsc.edu and https://brain-vasc.cells.ucsc.edu) as a pre-annotated Seurat object and used directly for Gene expression and/or Connectome analysis as described above.

We applied the **scDrugLink** method to predict drug sensitivity and inhibition effects across endothelial cell (EC) subclusters. This approach integrates drug target expression with transcriptomic reversal scores derived from CMAP L1000 perturbation signatures to compute drug-specific scores for each cell type. Mutant and control choroid EC clusters were excluded from the analysis to focus on brain-relevant populations. The angiogenic 2 cluster was compared to all EC clusters from *Alk1l/l* control mice. Drug sensitivity/resistance and inhibition/promotion scores were calculated and aggregated using the **scDrugLink** R package (https://github.com/LHBCB/scDrugLink). For each drug, the inhibition weight in a given cell type was computed by integrating **Cliff’s Delta** and the **adjusted p-value** derived from a within-cell-type **Wilcoxon rank-sum test** comparing mutant and control cells. Drugs with both high inhibition weights and high sensitivity scores were prioritized as repurposing candidates for targeting angiogenic 2 ECs.

### HUVECs culture

Cells were cultured in EBM2 basal medium supplemented with the SingleQuots Supplements. Cells were transfected with different siRNA (**Table 4**) using Lipofectamine RNAiMAX (Invitrogen, 13778150). 48h after transfection, cells were treated with BMP9 (R&D system, 3209-BP-010) at 5ng/mL during 24h.

**Table 4.**

| **siRNA** | **Company** | **References** |
| --- | --- | --- |
| Hu ON-TARGETplus Non-targeting Pool | Dharmacon | D-001810-10-05 |
| Hu *ACVRL1-6* | Qiagen | SI02758392 |
| Hu *ACVRL1-5* | Qiagen | SI02659972 |
| Hu *SMAD 1/4* | Qiagen | SI00020027 |
| Hu *ENG 6* | Qiagen | SI02663031 |

### mLECs isolation

CD31 antibody (BD Pharmingen, 550274) was incubated with dynabeads (ThermoFisher, 11035) at 4°C overnight and then washed with PBS-0,1%BSA using magnetic separator DynaMag^TM-15^ (ThermoFisher, 12301D) and resuspended in PBS-BSA 0,01%. Lungs were digested 1 hour at 37°C using Collagenase type I (Worthington, LS004196). Reaction was stopped with DMEM 5% SVF, filtered through a 70 µm cell strainer (Fisher, 087712). After one step of washing the pellet was resuspended in PBS-0,1% BSA and 100uL of beads coated with CD31 were added and incubated during 30min, at RT under agitation. Using the magnetic separator wash the cells with PBS and process to cell lysis.

### mBECs isolation

For cell culture PNVP was separated enzymatically from INVP as previously described. Myelin and debris were removed with DMEM-BSA 25% gradient, and after one step of wash with PBS, cells were seeded in EBM2 Basal medium (Lonza CC-3121) with SingleQuots Supplements (Lonza CC-4133) and 20% FBS (Foetal Bovin Serum, Thermofisher, 12484028) on plate coating with collagen I (Thermofisher, A1064401). Selection of EC was made with puromycin (Santa Cruz Biotechnology, SC-108071A) at 10µg/mL during two days, and after at 2µg/mL.

For RNA extraction, total brain was digested with collagenase/dispase (sigma, 11097113001) during 1 hour supplemented with DNASE I (50µg/mL, Sigma, DN25) and TCLK (0,147µg/mL, Tosyl-lysine-chloromethylketone Sigma T7254). Myelin and cell debris were removed as previous described and ECs was isolated using beads coated with CD31 and the magnetic separators.

### Quantitative real-time PCR

Total RNA was extracted using Monarch® Total RNA Miniprep Kit (NEB, T2010S), reverse transcription with iScript Reverse Transcription Supermix (1708841) and RT-qPCR with iQ SYBR Green Supermix (Biorad, 1708882) and primers (**Table 5**). Data were normalized with housekeeping genes and mRNA relative expression was calculated using the double CT method.

**Table 5.**
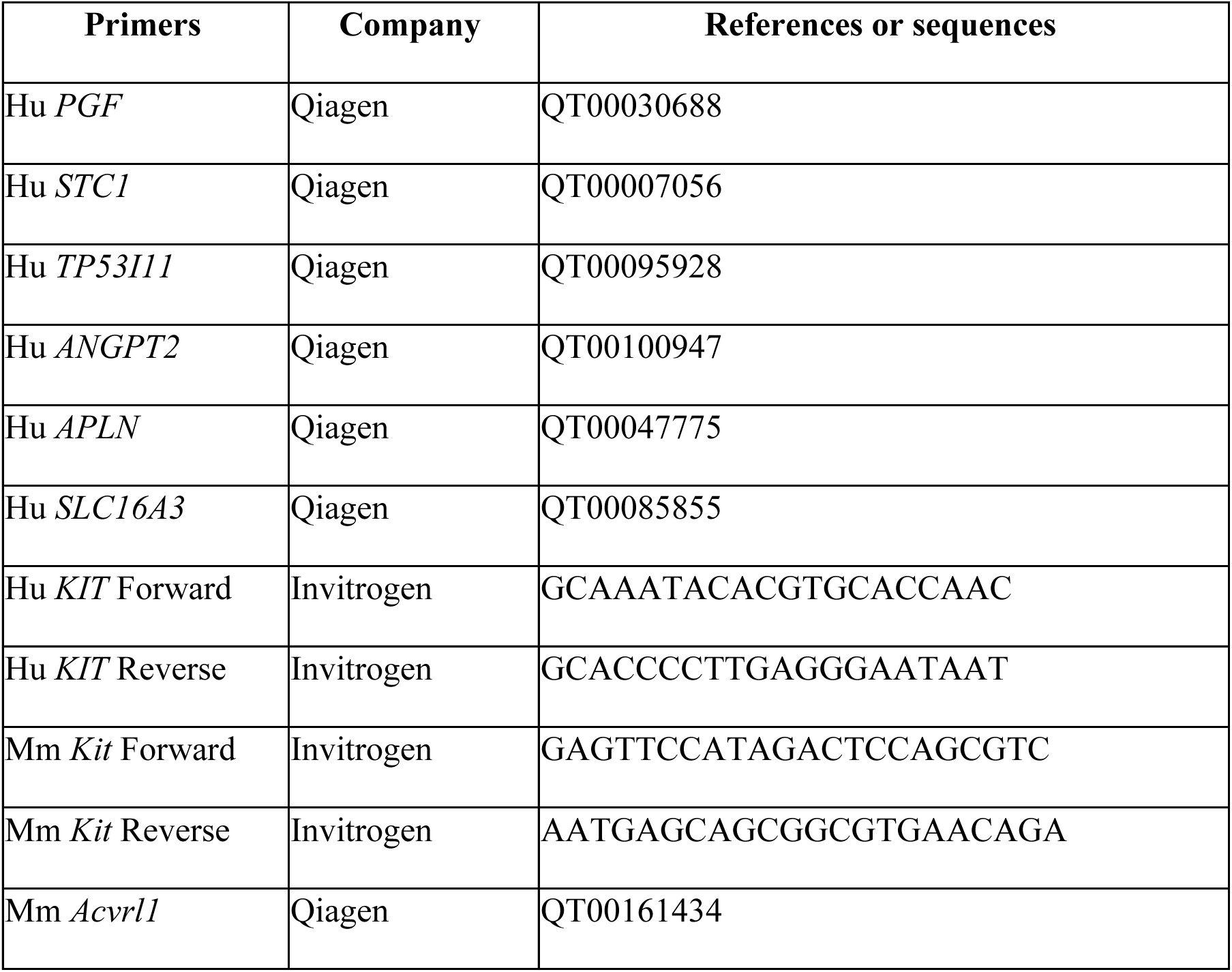

### Western blot

HUVECs were lysed with laemli’s buffer 4X (BioRad, 1610747) and protein lysate were loaded in NuPAGE™ MES SDS Running Buffer 20X (ThermoFisher, NP0002), transferred on nitrocellulose membrane (Bio-Rad, 1620112) and blocked 1 hour at RT on 3%BSA-TBST before the incubation with primary antibody (**Table 1**) at 4°C overnight. The day after membranes were washed with TBST, incubated 1 hour at RT horseradish peroxidase–labeled secondary antibodies (**Table 2**) and signal was revealed using SuperSignal™ West Femto Maximum Sensitivity Substrate (Thermo Fisher Scientific, 34096) or clarity western ECL substrate (Bio-Rad, 1705060S) on ChemiDoc MP imaging system (Bio-Rad). Analysis and quantification were made using Image Lab software (Bio-Rad).

### Statistical analysis

Statistical analysis and graph images were performed using GraphPad Prism 10 software. Comparisons across multiple groups were conducted using ordinary one-way ANOVA (for normally distributed data with equal variances), Welch’s ANOVA (for normally distributed data with unequal variances), or the nonparametric Kruskal–Wallis test (for non-normally distributed data). Statistical comparisons between two groups were conducted using either an unpaired t-test (for normally distributed data with equal variances), Welch’s t-test (for normally distributed data with unequal variances), or the Mann–Whitney test (for non-normally distributed data), as appropriate. P<0.05 was considered statistically significant. All results are represented as mean ± standard error of the mean (s.e.m).

### Materials availability

Materials are available upon reasonable request.

### Data and code availability

All data that support the findings of this study are available within the article and its supplemental information File and from the corresponding author upon reasonable request. The spatial and scRNAseq data discussed herein will be deposited in NCBI’s Gene Expression Omnibus (accession no. GSE297553). Code is available on github (https://github.com/gaelcge/brainAVM_Drape_scRNAseq).

## Results

### Brain endothelial *Alk1 d*eletion induces region-specific AVMs in the postnatal brain

To investigate bAVM formation in a model of HHT2, we used endothelial cell-specific and inducible *Alk1* knockout mice (*Alk1iECKO*). Tamoxifen was injected at P6, and the vascular phenotypes were assessed at P8 to circumvent early lethality observed in this model^26^ (**Fig. 1A**). We analyzed the perineural (PNVP) and intraneural (INVP) as distinct vascular compartments due to their different vessel compositions and hemodynamic environments^29^ (**Fig. 1A**). The PNVP, located at the brain surface, is primarily composed of veins, venules, and arteries and undergoes active remodeling during early postnatal development. It exhibits higher flow patterns compared to the INVP, which is largely composed of capillary beds within the brain parenchyma. Given that AVMs arise from aberrant arteriovenous (AV) connections and are strongly influenced by flow, we hypothesized that the PNVP would be more susceptible to bAVM formation upon loss of ALK1 signaling.

To examine the vascular network, we performed immunostaining for the endothelial marker CD31 on whole-mount brain preparations to assess the PNVP, and on coronal brain sections to examine the INVP (**Fig. 1A, Supp Fig. 1A**). To distinguish arteries within the vascular network, we co-stained with an antibody against α-smooth muscle actin (α-SMA), a marker of vascular smooth muscle cells^30^ (**Fig. 1A**). We observed that loss of ALK1 signaling induced vascular malformations in the PNVP compared to control littermates at P8 **(Fig. 1B**). Specifically, we detected a significant increase in the overall vascular area (**Fig. 1C**). This change is associated with a significant increase in the diameter of arteries, veins, and venules (**Fig. 1D**), along with a marked reduction in the proportion of CD31+ vessels smaller than 10 μm and a concomitant increase in vessels larger than 20 μm (**Fig. 1E**).

In line with previous reports that identified abnormal vein filling with blue latex in the presence of AV shunts, we also observed direct AV connections in the mutant PNVP, indicative of early bAVM formation (**Fig. 1F, G**). While we confirmed efficient *Alk1* deletion in ECs isolated from both the PNVP and INVP compartments (**Supp. Fig. 1B-C**), we did not observe significant vascular alterations in the INVP. Specifically, vessel diameter, vascular area, and direct AV connections remained unchanged in the mutant INVP compared to controls (**Fig. 1H-J**). These results indicate that a 48-hour loss of endothelial ALK1 signaling predominantly induces vascular malformations and AVMs in the PNVP.

### Single-Cell RNA sequencing reveals that *Alk1* deficiency induces EC transcriptomic aberrancy in PNVP and INVP ECs

To investigate the molecular mechanisms driving AVM formation in the PNVP, we performed single-cell RNA sequencing (scRNA-seq) of ECs isolated from both PNVP and INVP regions of the same brains in P8 *Alk1iECKO* and control mice (**Fig. 2A, supp fig. 2A**). We sequenced 1,195 *Alk1 l/l* and 2,193 *Alk1iECKO* PNVP ECs and 3,221 *Alk1 l/l* and 1,904 *Alk1iECKO* INVP ECs, with an average of 4,097 genes per cell.

**Figure 2:**
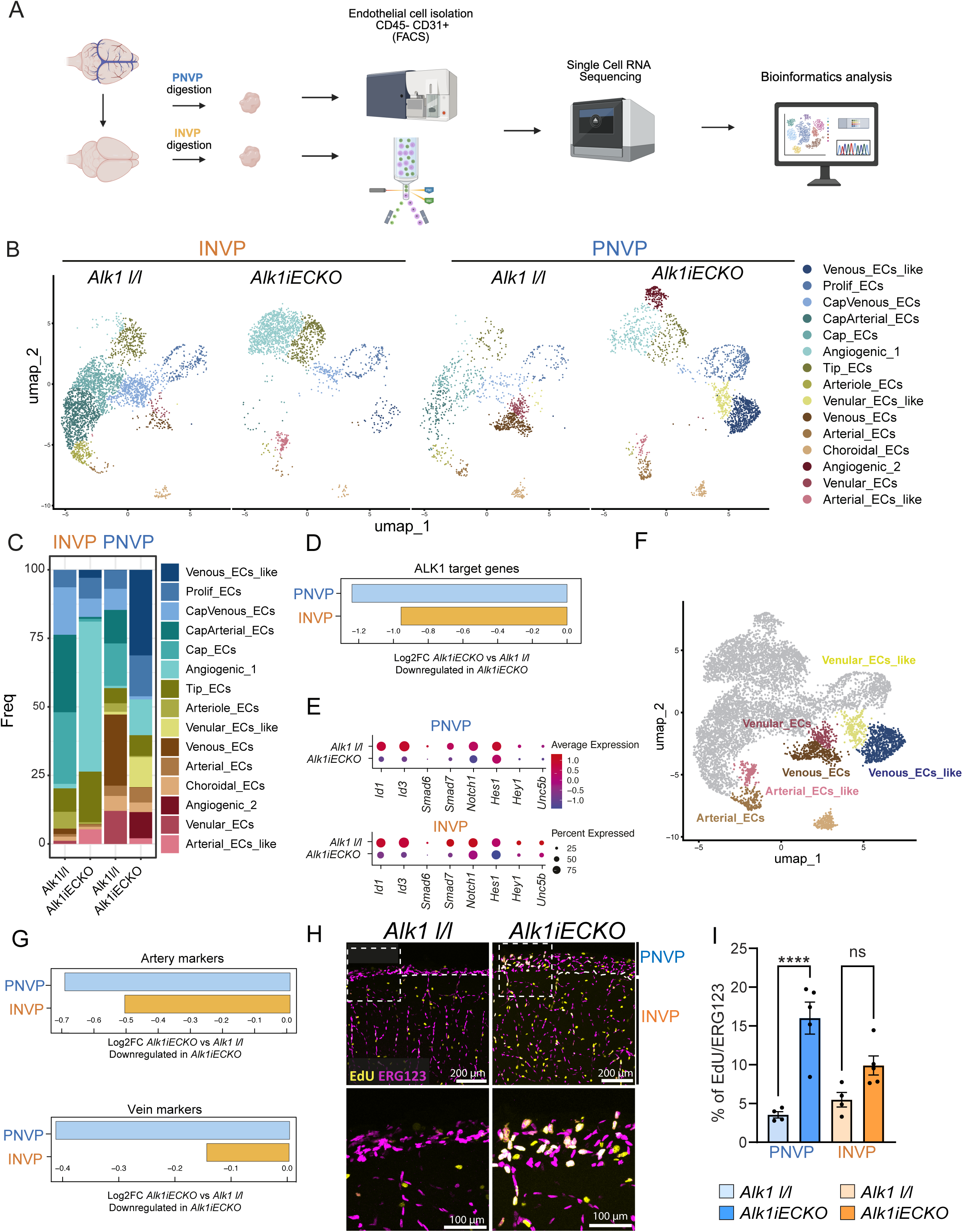
Single-cell RNA sequencing reveals distinct endothelial cell reprogramming in *Alk1iECKO* brains. **A**, Experimental workflow for scRNAseq of ECs from PNVP and INVP of *Alk1iECKO* and control mice at P8. **B**, UMAP plots of EC clusters from INVP and PNVP of *Alk1iECKO* and control brains. **C**, Proportion of EC clusters in indicated conditions. **D**, Barplot of VISION score log2FC (*Alk1iECKO* vs control) of ALK1 signaling target gene set in PNVP and INVP endothelial cells, highlighting loss of ALK1 signaling. **E**, Dot plot of the LK1 signaling target genes expression. **F**, UMAP plot of EC clusters from INVP and PNVP of *Alk1iECKO* and control brains highlighting arterial and venous ECs. **G-H**, Barplot of VISION score log2FC (*Alk1iECKO* vs control) of arterial (**G**) and venous (**H**) gene set signatures in PNVP and INVP ECs **I**, Double ERG and EdU immunolabelling of P8 *Alk1iECKO* and control brains. The white dotted line delineates the boundary between the PNVP and the INVP. **J**, Quantification of ERG+EdU+/ERG+ cells. Each dot represents one mouse. Error bars represent means ± s.e.m, *\*\*\*\*P<0.0001,* Ordinary one-way ANOVA (J) was performed.

UMAP visualization and unsupervised clustering revealed distinct EC subpopulations in both genotypes and brain regions (**Fig. 2B, supp fig. 2A-B**). In the INVP of control mice, we identified a typical continuum of arterial to venous ECs, with a predominance of capillary ECs, as well as proliferative and tip cell populations reflecting the active angiogenic phase of postnatal development. In contrast, the PNVP of control brains was mostly composed of venous, venular, and arterial ECs, consistent with a vascular bed undergoing hemodynamic remodeling at this stage (**Fig. 2C**). Moreover, in the PNVP dataset we also detected capillary and tip ECs resembling those in the INVP, which could reflect partial contamination due to enzymatic overdigestion of the cortical INVP during PNVP ECs isolation. Nonetheless, the major EC populations identified across samples were consistent with previously described vascular zones and cell identities^31^.

To confirm efficient deletion of *Alk1*, we performed pathway activity score analysis (VISION package) using a custom ALK1 signaling signature geneset^32^ **(Table 6**). This revealed a marked reduction in ALK1 target gene expression across all mutant EC clusters, in both PNVP and INVP (**Fig. 2D, E**). In the PNVP, *Alk1iECKO* ECs showed a striking loss of arterial, venular, and venous identity, accompanied by the emergence of new arterial-like, venular-like, and venous-like clusters (**Fig. 2F, G, supp fig. 2C, D**). Similar, though less pronounced, shifts in AV EC identities were also observed in the INVP. Notably, we observed that *Alk1* deletion led to increased EC proliferation predominantly in the PNVP, consistent with the vascular malformations seen *in vivo* (**Fig.1 B-J, Fig2. B-C**). This was supported by EdU/ERG co-labeling experiments, which showed a significant increase in proliferating ECs specifically in the mutant PNVP (**Fig. 2 H, I**).

Given prior studies suggesting that blood–brain barrier (BBB) defects in AVMs may contribute to neurological symptoms^33–35^, we also assessed BBB-related gene expression. Using a custom BBB geneset (**Table 6**), pathway activity analysis indicated that while the INVP normally displays a stronger BBB signature than the PNVP, loss of ALK1 signaling resulted in a more pronounced BBB gene downregulation in the PNVP (**Supp Fig. 2 E, F**), highlighting region-specific vascular vulnerabilities in P8 *Alk1iECKO* brain.

### Characterization of PNVP-enriched angiogenic EC clusters in *Alk1iECKO* Brains

BMP9/ALK1 signaling is a key regulator of angiogenic sprouting and tip cell specification^32^. Consistent with previous findings in the retina, *Alk1* deletion resulted in an increased number of tip ECs in the brain (**Fig. 3A-C**). In addition, it induced the emergence of two new angiogenic endothelial clusters in both the INVP and PNVP regions (**Fig. 2B, C, Fig. 3A**), annotated as angiogenic 1 and angiogenic 2 clusters based on the expression of tip EC markers (**Fig. 3D**), consistent with the heightened angiogenic activity characteristic of AVMs^36^.

**Figure 3:**
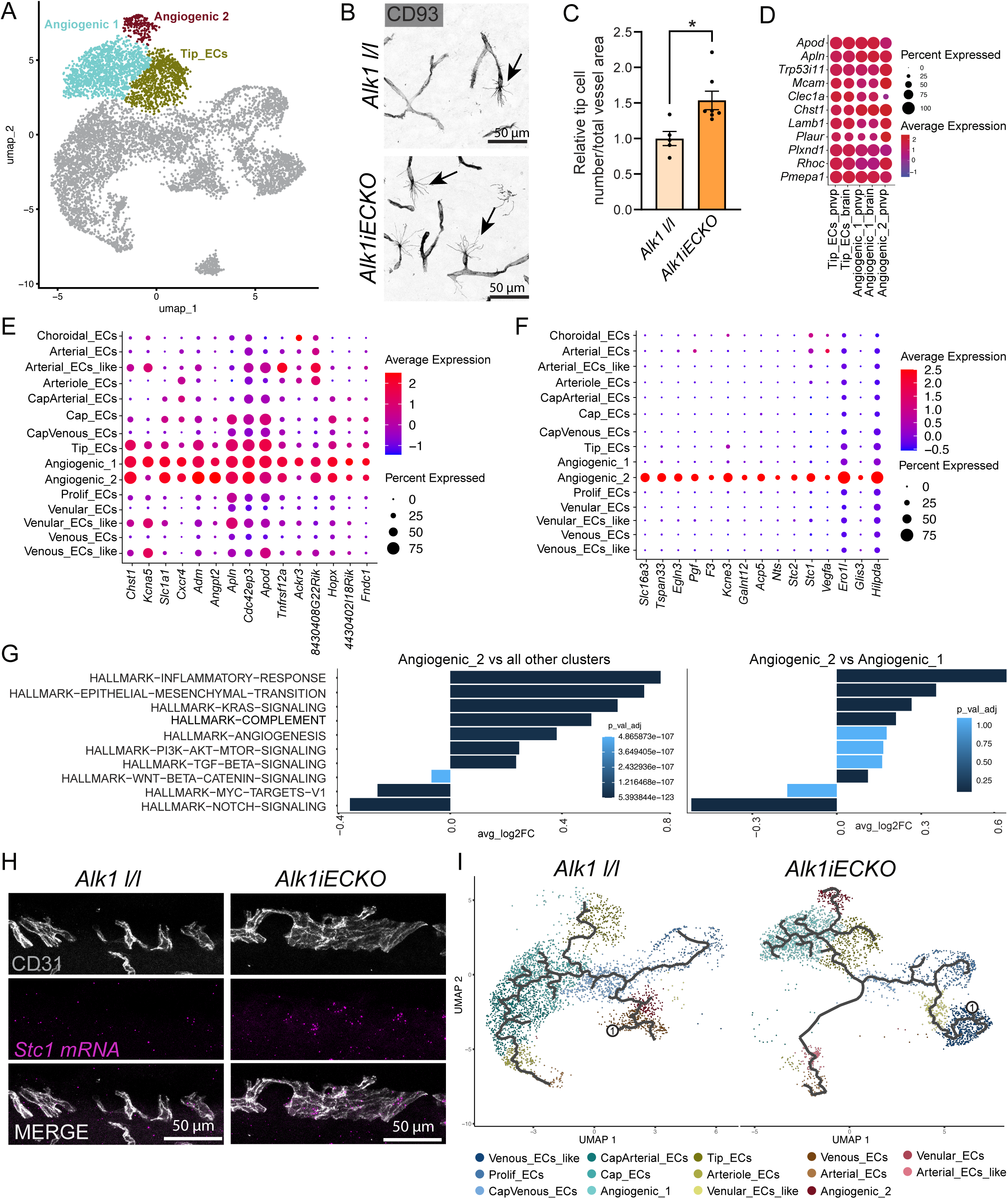
Characterization of PNVP-enriched angiogenic EC clusters in Alk1iECKO brains. **A**, UMAP plots of EC clusters from INVP and PNVP of *Alk1iECKO* and control brains highlighting angiogenic clusters. **B,** CD93 immunostaining showing an increase of tip cell number in the INVP of *Alk1iECKO* mice. **C**, Quantification of tip cell numbers in Alk1iECKO and control INVP. **D**, Dot plot of tip cell markers expression. **E-F**, Dot plot of top 15 angiogenic 1 (E) and angiogenic 2 (F) cluster markers. **G**, Barplot of VISION score log2FC in angiogenic 2 ECs against all other clusters for the top 10 most significant pathways from the hallmark MsigDB genesets. **H**, *In situ* hybridization for *Stc1* mRNA confirming increased expression in PNVP endothelial cells. **I**, Trajectory analysis of endothelial cells from control and *Alk1iECKO* mice at P8. Each dot represents one mouse. Error bars represent means ± s.e.m, *\*P<0.05,* Mann-Whitney test (C) was performed.

While the angiogenic 1 ECs exhibited strong transcriptomic similarity to canonical tip cells, angiogenic 2 ECs displayed a divergent expression profile marked by genes such as *Slc16a3*, *Pgf*, and *Stc1* (**Fig. 3E, F**). Differential pathway activity analysis revealed that angiogenic 1 ECs displayed upregulation of pathways involved in cardiac contraction and regulation of blood circulation, whereas angiogenic 2 ECs were enriched in pathways related to glycolysis and hypoxia (**Supp. Fig. 3A, B**). Focusing on the hallmark pathways from MsigDB, the angiogenic 2 cluster also exhibited increased enrichment for several brain AVM-related gene signatures compared to other EC clusters, and particularly relative to the angiogenic 1 cluster (**Fig. 3G**). These included pathways previously reported in human bAVMs, such as inflammatory response, epithelial-to-mesenchymal transition (EMT), complement, angiogenesis, and PI3K/mTOR signaling^24^. We also noted dysregulation of KRAS and NOTCH signaling, both implicated in AVM pathogenesis^20^. Moreover, angiogenic 2 ECs expressed genes previously reported to be upregulated in *Alk1*-deficient ECs, such as *Itgb1* and *Itgav*^18^, and genes recently identified as candidate drivers in HHT, including *Jak2*, *Angptl4*, and *Foxo1*^37^ (**Supp. Fig. 3C**). These data point to functionally divergent angiogenic states shaped by tissue context: angiogenic 1 ECs predominated in the INVP, replacing capillary ECs, whereas angiogenic 2 ECs were selectively enriched in the PNVP.

Given that AVMs predominantly form in the PNVP at P8, we hypothesized that the angiogenic 2 represent ECs within AVMs. Supporting this, *in situ* hybridization (ISH) for *Stc1* confirmed increased expression in PNVP ECs of *Alk1iECKO* mice (**Fig. 3H, Supp. Fig. 3D**), consistent with angiogenic 2 ECs comprising the AVM endothelium. Because AVMs are believed to originate from the venous vascular bed^18,19^, we performed

RNA trajectory analysis using the vein cluster as root, allowing pseudotime inference based on transcriptomic variations across cells which can then be visualized on UMAP dimensions. In control brains, venous ECs transitioned toward proliferative or capillary ECs, which then gave rise to arterial ECs, consistent with known postnatal vascular maturation (**Fig. 3I**). In contrast, in *Alk1iECKO* ECs, venous ECs progressed to angiogenic 1 ECs, and subsequently gave rise to tip cells and angiogenic 2 ECs (**Fig. 3I**).

These data support a venous origin for AVMs and suggest that angiogenic 1 ECs represent an early reprogrammed state that can evolve into angiogenic 2 ECs. Altogether, our results point to angiogenic 2 ECs as a transcriptomic signature of AVM-forming endothelium.

### Mouse *Alk1iECKO* Angiogenic_2 and human AVM endothelial clusters share AVM molecular markers including KIT

To explore the transcriptional overlap between the angiogenic 2 cluster in *Alk1iECKO* brains and endothelial clusters identified in human brain AVMs by Winkler E. et al.^24^, we next examined shared molecular markers. Cross-species comparison revealed common upregulated genes involved in angiogenesis, including *PGF*, *STC2*, *APLN*, and *KIT* (**Fig. 4-C, Supp. Fig. 4A, B**). We also found these markers enriched in other human brain dataset published by Wälchli T. et al^25^ (**Supp. Fig.4C**), suggesting conserved AVM-associated endothelial reprogramming.

**Figure 4:**
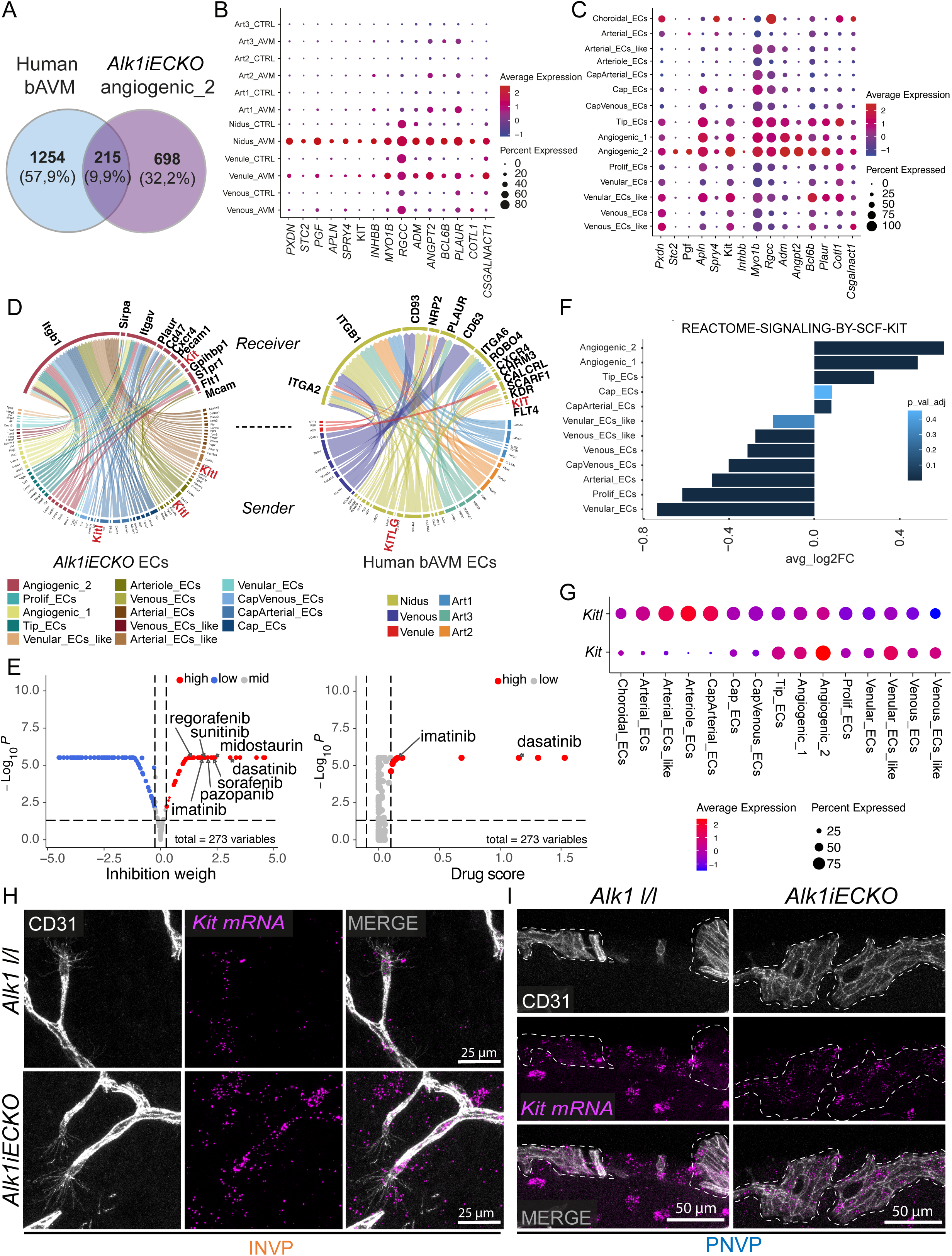
Mouse Alk1iECKO Angiogenic_2 and human AVM endothelial clusters share AVM molecular markers including KIT. **A**, Venn diagram of marker genes (log2 FC 0.25, adjusted p-value 0.05) from human brain nidus and *Alk1iECKO* angiogenic 2 cluster. **B-C**, Dot plot of top 10 common markers expression in human (B) and mouse (C) ECs. **D**, Circos plot representation of enriched ligand-receptor interactions in mouse (min.pct 0.5, min.z 0.5) and human (min.pct 0.2, min.z 0.5) EC clusters. **E**, Drug repurposing analysis applied to the *Alk1iECKO* angiogenic 2 EC cluster. Left: Network-based inhibition weight of angiogenic 2 cluster compared to all control EC clusters. Right: Top candidate drugs ranked by predicted drug score. **F,** Barplot of VISION score log2FC of KIT signaling between mouse EC clusters. **G**, Dot plot of *Kit* and *Kitl* expression. **H-I**, *In situ* hybridization for *Kit* mRNA confirming increased expression in INVP tip cells and PNVP endothelial cells.

To characterize the vascular communication profile of these AVM-associated clusters, we performed ligand–receptor connectome analysis using the Connectome R package^38^, examining signaling interactions of the *Alk1iECKO* angiogenic 2 cluster and the human AVM nidus cluster with other EC populations (**Fig. 4D, Supp. Fig.4D**). Both clusters exhibited widespread interactions with other EC types, primarily through angiogenic, integrin, and ECM-related signals, such as *Itgb1*, *Pgf*, and *Kit* (**Fig. 4D, Supp. Fig.4D**). Surprisingly, we detected very few ligand–receptor interactions between angiogenic 2 and proliferative EC clusters, despite both being present in the mouse AVM dataset (although proliferative clusters were not detected in the human AVM dataset).

Next, to further prioritize therapeutic targets, we leveraged a single-cell drug repurposing approach (scDrugLink), which integrates gene expression, drug targets, and perturbation signatures. This analysis revealed that angiogenic 2 ECs displayed high inhibition weights and drug sensitivity scores for several receptor tyrosine kinase (RTK) inhibitors, including KIT inhibitors such as dasatinib and imatinib (**Fig. 4E, Table 7, Table 8**). Taken together, the prominence of *Kit* among the shared angiogenic genes, its predicted ligand–receptor interactions, and its prioritization through computational drug screening collectively highlight KIT as a compelling and targetable vulnerability of the angiogenic 2 state. *Kit* encodes a tyrosine kinase receptor activated by its ligand, stem cell factor (SCF, encoded by *Kitl*), which regulates cell migration and proliferation in immune and cancer contexts^39,40^. KIT is also an attractive candidate for therapeutic targeting, as KIT inhibitors are widely used in oncology, suggesting potential for drug repurposing if KIT signaling contributes to brain AVM pathogenesis^41^. Pathway activity analysis revealed robust enrichment of KIT signaling in the angiogenic 2 cluster, with a lower increase in angiogenic 1, consistent with a potential progression from angiogenic 1 to angiogenic 2 ECs (**Fig. 4F**). In our dataset, *Kitl* was broadly expressed across EC clusters, with highest levels in arterial ECs and lowest in venous ECs, while Kit showed an inverse gradient, low in venous ECs and absent in arterial ECs (**Fig. 4G**). In control ECs, Kit expression was largely restricted to tip cells, whereas in mutant ECs it was broadly upregulated in angiogenic EC populations, with predominant enrichment in the angiogenic 2 cluster. Notably, both *Kit* and *Kitl* were co-expressed in angiogenic 2 ECs, supporting the possibility of cell-autonomous KIT activation, consistent with their predicted connectome interactions (**Fig. 4D-G**). We then validated the upregulation of *Kit* mRNA in *Alk1iECKO* ECs using *ISH*, which confirmed strong expression in INVP tip cells and a mosaic increase in a subset of PNVP ECs, consistent with its specific enrichment in the angiogenic 2 cluster (**Fig. 4H, I, Supp. Fig. 4E**). Interestingly, this upregulation appeared brain-specific, as *Kit* expression remained unchanged in lung ECs isolated from *Alk1iECKO* mice (**Supp. Fig. 4F**).

### ALK1 signaling suppresses KIT and Angiogenic_2 gene expression *in vitro*

We next investigated whether BMP9/ALK1 signaling directly regulates KIT expression in endothelial cells. HUVECs were transfected with either control or ALK1 (*Acvrl1*) siRNA, treated for 24 hours with 5 ng/mL BMP9, and analyzed for KIT expression at both the mRNA and protein levels using qPCR and western blotting, respectively. In control-transfected cells, KIT was expressed at high levels under basal conditions, but BMP9 treatment robustly induced phospho-SMAD1/5/9 and led to a strong reduction of KIT expression at both the protein and mRNA levels (**Fig. 5A-C**). Silencing ALK1 prevented BMP9-induced SMAD1/5/9 phosphorylation and rescued KIT expression, confirming the requirement of ALK1 for BMP9-mediated KIT repression (**Fig. 5A-C**). Moreover, knockdown of *SMAD4* significantly increased KIT levels as well (**Fig. 5D**). These in vitro findings are consistent with recent RNA-seq data showing that brain ECs from *Alk1* and *Smad4* mutant mice exhibit increased Kit expression^42^ (**Supp. Fig. 5A-B**).

**Figure 5:**
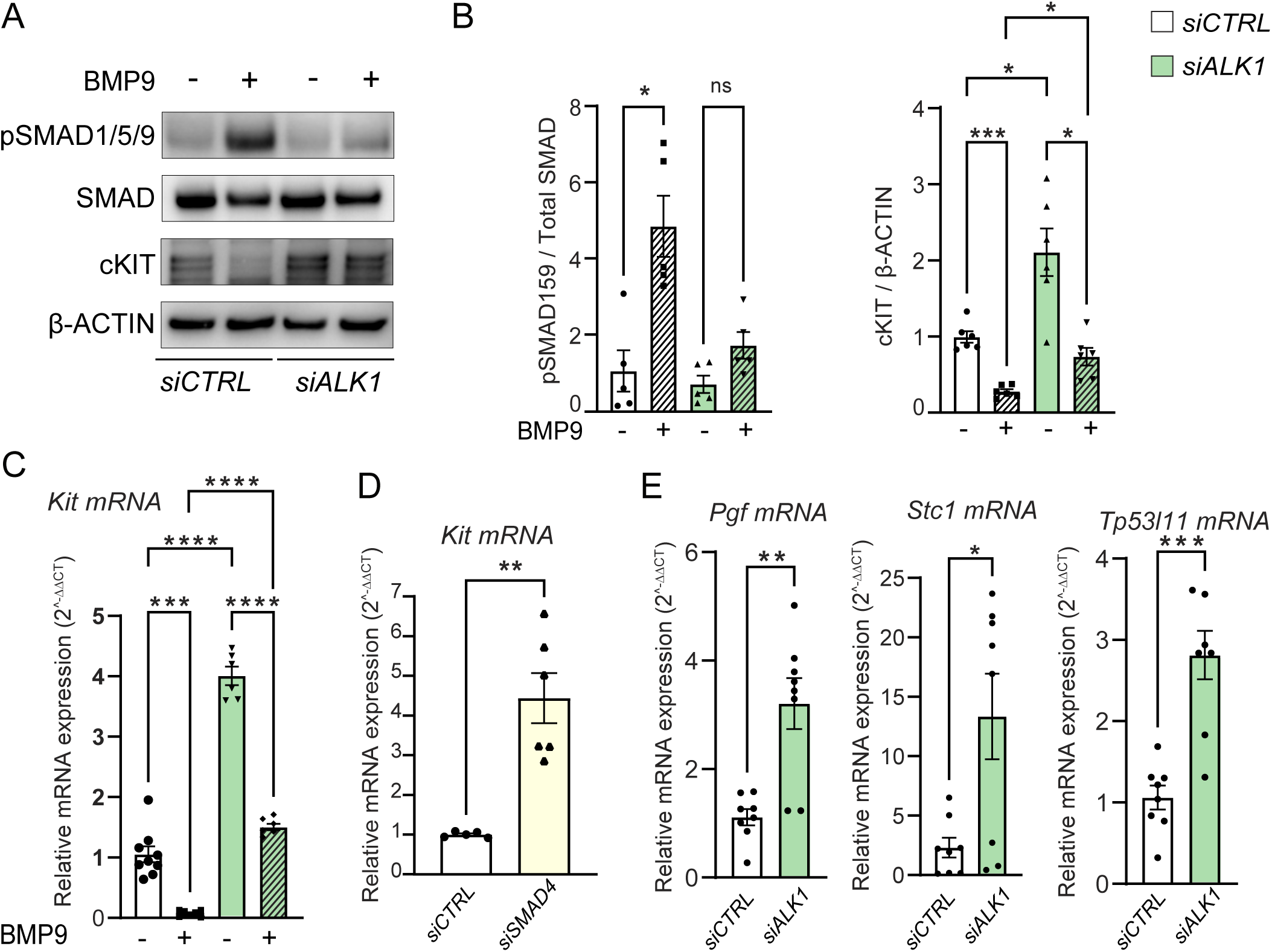
ALK1 signaling under suppresses KIT and Angiogenic_2 gene expression *in vitro*. **A**, Immunoblot detection of p-SMAD1/5/9, total SMAD1, KIT and β-Actin in HUVECs treated or not with BMP9 (5 ng/mL, 24H), and transfected with siRNA control or *ALK1*. **B**, Quantification of p-SMAD1/5/9/SMAD1 (left) and KIT/β-Actin (right) protein levels of immunoblots corresponding to A. **C-E**, qPCR analysis of *KIT* expression in HUVECs treated with *ALK1* (C), or *SMAD4* (D) siRNA compared to control. **E**, qPCR analysis of selected angiogenic 2 markers in HUVECs treated with *ALK1*siRNA compared to control. Each dot represents one experiment. Error bars represent means ± s.e.m, *\*P<0.05, **P<0.01, ***P<0.001, ****P<0.0001,* Brown-Forsythe and Welch ANOVA (B,C), Welch’s t test (D0, Unpaired t test or Mann-Whitney test (E) were performed.

In addition to KIT, BMP9/ALK1 signaling also repressed the expression of several angiogenic 2 markers *in vitro*, including *PGF* and *STC1* (**Fig. 5E, Supp. Fig. 5C**). Altogether, our data indicate that BMP9/ALK1/SMAD4 signaling directly regulates the angiogenic 2 transcriptional program and KIT expression in ECs, rather than acting through secondary changes in cell identity associated with malformation. These findings suggest aberrant KIT signaling may contribute to the initiation of brain AVM pathogenesis in *Alk1iECKO* mice.

### Pharmacological KIT inhibition prevents bAVM formation in *Alk1iECKO* Mice

To investigate the functional role of KIT signaling in the initiation of brain AVMs, we treated *Alk1iECKO* and control littermate mice with KIT inhibitors^41,43,44^ or vehicle controls at the time of tamoxifen administration, and analyzed the PNVP vascular network at the peak of AVM formation (**Fig. 6A**). Masitinib, a selective RTK inhibitor with its lowest IC₅₀ against KIT, significantly reduced veins and venules diameter and prevented vascular malformations induced by ALK1 signaling loss (**Fig 6B, C**). To further validate these findings, we administered a selective KIT-blocking antibody to *Alk1iECKO* mice, which improved venous diameter (**Fig. 6D-E**). Finally, to assess the potential therapeutic impact of KIT inhibition, mice were injected with Imatinib (or Gleevec), an FDA-approved multi-kinase inhibitor with activity against KIT. Consistent with our scDrugLink analysis (**Fig. 4E**), which identified Imatinib among the top compounds predicted to inhibit angiogenic 2 ECs, Imatinib treatment markedly improved PNVP vascular architecture **(Fig. 6F, G**). Importantly, these drugs had no detectable impact on PNVP vasculature of control mice (**Supp. Fig. 6A-D**).

**Figure 6:**
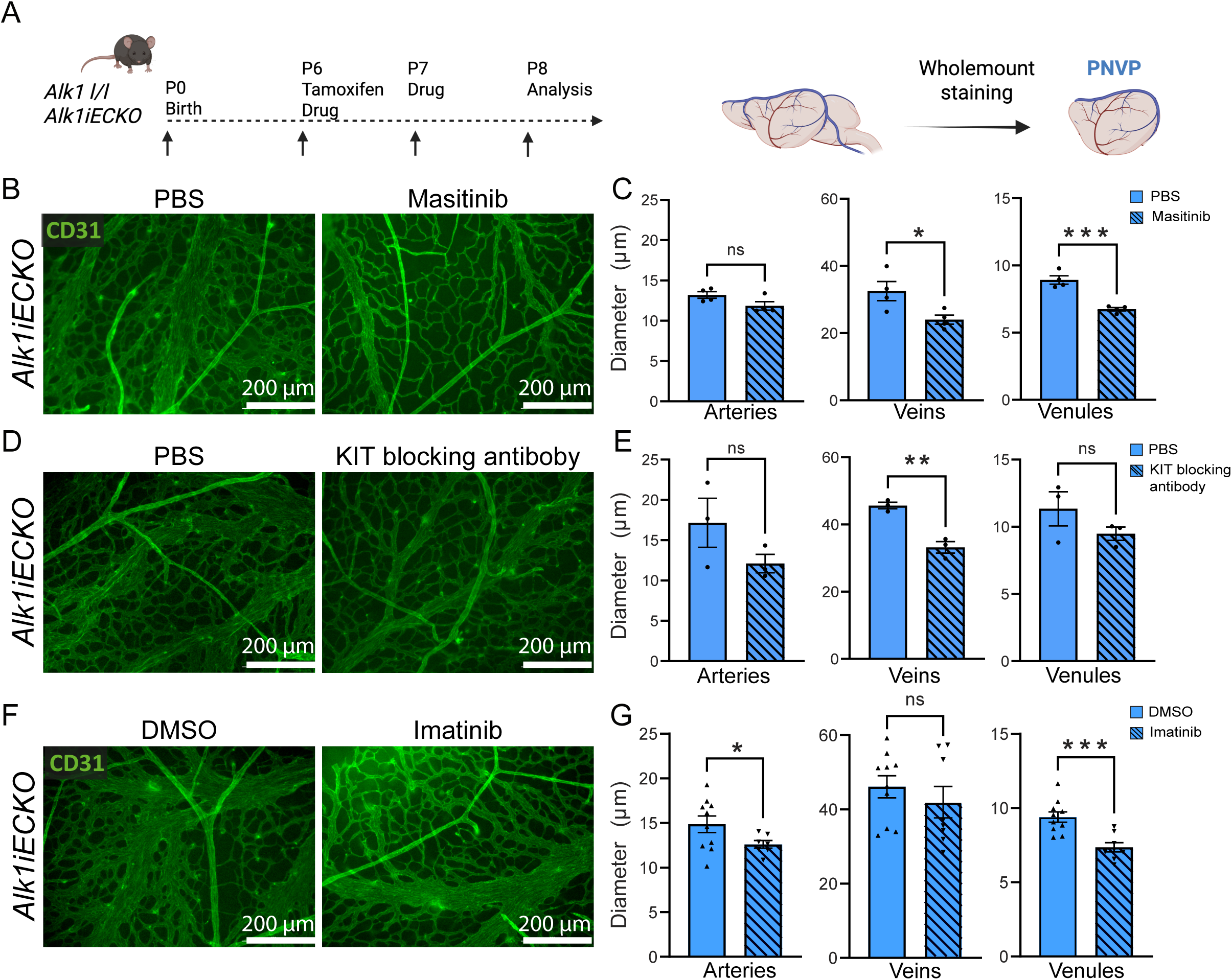
Pharmacological KIT inhibition prevents bAVM formation in *Alk1iECKO* Mice. **A**, Experimental timeline of tamoxifen and KIT inhibitor administration in *Alk1iECKO* and control mice. **B, D, F**, CD31 immunostaining of the PNVP of P8 *Alk1iECKO* mice injected with imatinib (**B**), masitinib (**D**), and KIT blocking antibody (**F**), along with their corresponding vehicle controls. **C, E, G**, Quantification of vessel diameter in the PNVP of P8 *Alk1iECKO* mice injected with the indicated inhibitor. Each dot represents one mouse. Error bars represent means ± s.e.m, *\*P<0.05, **P<0.01, ***P<0.001,* Mann-Whitney test, Welch’s t test or Unpaired t test (B, D, F) were performed.

## Discussion

Vascular heterogeneity has emerged as a defining feature of brain AVMs, yet the transcriptional programs underlying lesion formation remain poorly understood. Here, we used single-cell transcriptomics to characterize EC heterogeneity and transcriptional reprogramming in a mouse model of HHT2 bAVMs. We found that AVMs form at the brain surface (PNVP) within 48 hours of *Alk1* inactivation, driven by the emergence of a distinct angiogenic EC cluster enriched for *Kit* expression. We further demonstrated that BMP9/ALK1/SMAD4 signaling regulates *Kit* transcription in brain ECs. Finally, we showed that pharmacological inhibition of KIT signaling prevents bAVM onset in this model **(Supp. Fig. 6E**).

Our data reveal that the PNVP is more susceptible to AVM formation than the INVP, likely due to its arteriovenous architecture and associated hemodynamic forces. This regional vulnerability is consistent with the venous origin of AVMs and their predilection for high-flow regions observed in both mouse models and human brain pathology^35,45^. Trajectory analysis further supports this idea, showing that the angiogenic and proliferative EC clusters driving AVM formation originate from venous ECs. Notably, AVMs emerge in an angiogenic context, yet the PNVP at this developmental stage is undergoing physiological regression, in contrast to the actively expanding INVP^29^. It is therefore surprising that loss of ALK1 signaling not only prevents PNVP regression but also induces endothelial proliferation and AVM formation within this normally regressing vascular bed. The molecular mechanisms governing PNVP regression during postnatal development remain poorly understood; however, our findings point to ALK1 as a critical regulator of this process.

Moreover, the apparent confinement of AVMs to the PNVP at P8 may reflect a temporal constraint, and in models with longer survival, these malformations are likely to extend into the INVP. In patients, brain AVMs are most frequently located in superficial cortical regions but can also occur in deeper structures, including the basal ganglia and thalamus^46^. These findings underscore the importance of regional vascular context in AVM pathogenesis. Identifying such regional phenotypic differences provides a framework to better design studies and could facilitate the discovery of new molecular drivers involved in brain AVM development.

Using an original regional PNVP and INVP EC isolation strategy, our study reveals a transcriptional reprogramming of AVM ECs characterized by both proliferative and angiogenic clusters. Aberrant EC proliferation is not merely a consequence of AVMs but plays a key role in their initiation and growth. Genet et al. showed that CDK (cyclin-dependent kinase) 4/6 inhibitor (CDK4/6i) palbociclib treatment reduces retinal *Alk1iECKO* AVM by limiting EC proliferation^26^.

BMP9/ALK1 signaling plays a central role in regulating angiogenic sprouting and tip cell specification ^32^, and accordingly, *Alk1* deletion in the brain induced the emergence of two transcriptionally distinct angiogenic endothelial clusters. Angiogenic 1 ECs, enriched in the INVP, displayed strong transcriptomic similarity to canonical tip cells and appeared to emerge early in the AVM trajectory. RNA velocity analysis suggests that angiogenic 1 ECs may give rise either to sprouting tip cells or to angiogenic 2 ECs associated with AVM formation. The molecular mechanisms governing this fate decision remain unknown but may depend on contextual cues such as hemodynamic forces. As AVMs arise in high-flow environments, elevated shear stress may bias angiogenic 1 ECs toward an angiogenic 2 state, thereby promoting AVM onset.

We identified an angiogenic 2 population characterized by both angiogenic and inflammatory signatures, as previously described by Winkler et al. in human bAVM samples^24^. Notably, while our data are derived from a mouse model of HHT2 and theirs from human non-HHT AVMs, the convergence of angiogenic and inflammatory pathways in both contexts suggests conserved molecular mechanisms across species and AVM etiologies. This is particularly striking given the distinct developmental origins and genetic backgrounds involved. Interestingly, established human brain AVMs did not exhibit a proliferative EC cluster, suggesting that the angiogenic EC cluster may represent a more relevant target for treating established AVMs.

We found the angiogenic 2 cluster is enriched for *Kit* expression. Interestingly, while *Kit* is mostly confined to tip cells in control mice, it is more broadly expressed across several mutant EC clusters, albeit at a lower level than the angiogenic 2 cluster. This suggests that *Kit* is a robust marker of angiogenic EC state, although it is not restricted to tip cells in pathological angiogenesis such as AVMs. Moreover, *SMAD4* knockdown significantly increased KIT levels, consistent with RNA-seq data showing increased *Kit* expression in *Alk1* and *Smad4* mutant brain ECs^42^. Therefore, ALK1–SMAD4 signaling negatively regulates *Kit* and that this repression is uncoupled from ENG. These findings suggest that *Kit* upregulation is a feature of ALK1/SMAD4-deficient AVMs, but may not be a marker of HHT1-associated lesions.

To prioritize potential therapeutic targets, we applied a single-cell drug repurposing analysis. This approach revealed that the angiogenic 2 cluster displayed high inhibition weights and sensitivity scores for several KIT-targeting agents such as imatinib and dasatinib. Pharmacological inhibition of KIT signaling efficiently prevented AVM, underscoring its functional relevance. The role of KIT in tip cell and AVM remains to be fully elucidated. In addition to its well-known function in angiogenesis^47–49^, KIT signaling also promotes PI3K/MTOR signaling^50^ and HIF1a expression ^51^, both important regulators of glycolysis^52,53^, which we found strongly upregulated in angiogenic 2 ECs. KIT may also influence key processes involved in AVM pathophysiology, including EC migration, polarity, and flow-responsive PI3K signaling^21^. Therefore, future studies should evaluate how KIT signaling interacts with other pathways involved in AVM formation.

Moreover, since Kit expression is not entirely restricted to the angiogenic 2 cluster, it is possible that its inhibition affected multiple cell states. Future studies using more specific genetic tools will be necessary to dissect the precise contribution of angiogenic 2 ECs and to validate additional markers more selectively enriched in this cluster, such as *Pgf*. For example, recent studies have demonstrated that targeting ANGPT2, a marker more specifically enriched in the angiogenic 2 cluster, efficiently rescues AVMs in the postnatal retina and brain^42^.

To our knowledge, this is the first identification of an AVM-specific EC cluster in an HHT mouse model using scRNA-seq. While previous single-cell studies, including those by Genet et al. in the retina^26^, did not resolve a distinct AVM EC population, we were able to validate several of our AVM markers in their dataset. This discrepancy may stem from regional differences in vascular maturation and heterogeneity. Compared to postnatal retina, the brain vasculature, with its advanced maturation, may provide a more sensitive system for detecting transcriptomic reprogramming in AVM models, highlighting its value for future mechanistic studies. Therefore, our study offers a valuable resource for uncovering novel molecular targets and developing new therapeutic strategies to treat bAVMs.

### Limitations

Our study has several limitations. First, while our model recapitulates key aspects of HHT-associated brain AVMs, most human AVMs are sporadic and not linked to HHT, which may limit the generalizability of our findings. Second, we identified EC transcriptomic reprogramming at an early stage of brain AVM formation, restricting our ability to capture dynamic changes in EC states. A longitudinal analysis at later stages will be critical to define the ontogeny of AVM EC populations and to identify progressive reprogramming events that could contribute to neurovascular complications. Third, we did not validate *Kit* expression in human HHT2 brain AVMs, which limits the direct translational relevance of our findings. Fourth, although KIT signaling inhibition prevented AVM onset, we did not assess its efficacy in reversing or improving established lesions, a crucial distinction for clinical translation, given that AVMs are typically diagnosed after lesion formation. Addressing these limitations will be essential for refining our understanding of AVM initiation and progression and for developing targeted therapeutic strategies.

## Abbreviations

ALK1: Activin Receptor-like Kinase 1
bAVMs: brain Arteriovenous Malformations
BBB: Blood Brain Barrier
BMP: Bone Morphogenetic Proteins
ECs: Endothelial Cells
HHT: Hereditary Hemorrhagic Telangiectasia
HUVEC: Human Umbilical Venous Endothelial Cell
INVP: Intraneural vascular plexus
mBEC: mouse Brain Endothelial Cell
mLEC: Mouse Lung Endothelial Cell
PNVP: Perineural vascular plexus
SCF: Stem Cell factor
scRNAseq: Single-cell RNA sequencing
siRNA: small interfering RNA

## Acknowledgments

We would like to thank Dr. Typhaine Anquetil for her invaluable help with the scRNA-seq experiments. We also extend our gratitude to Dr. Faughnan Marie and Dr. Sabine Bailly for their insightful discussions and contributions to the project. Our sincere thanks to the animal facility for their dedicated care of the animals used in this study.

---

**Supp. Figure 1.**
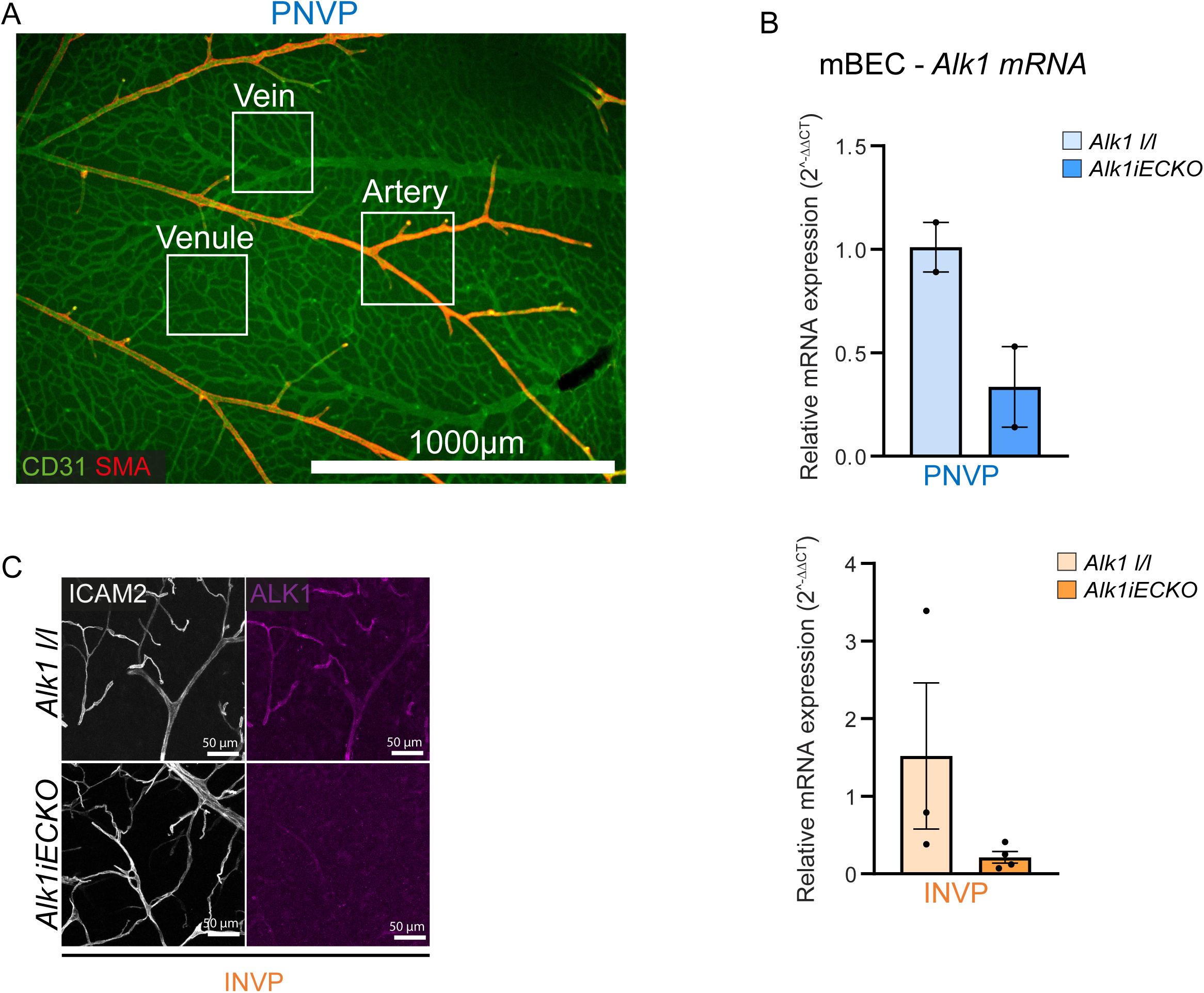
**A**, Representative images showing CD31 and αSMA immunostaining and wholemount imaging of the PNVP at P8. White squares indicate representative areas used for quantification. **B**, qPCR analysis of *Alk1* expression in isolated brain ECs from PNVP and INVP *Alk1iECKO* and control mice. **C**, CD31 and ALK1 immunostaining of the INVP of P8 *Alk1iECKO* and control mice.

**Supp. Figure 2.**
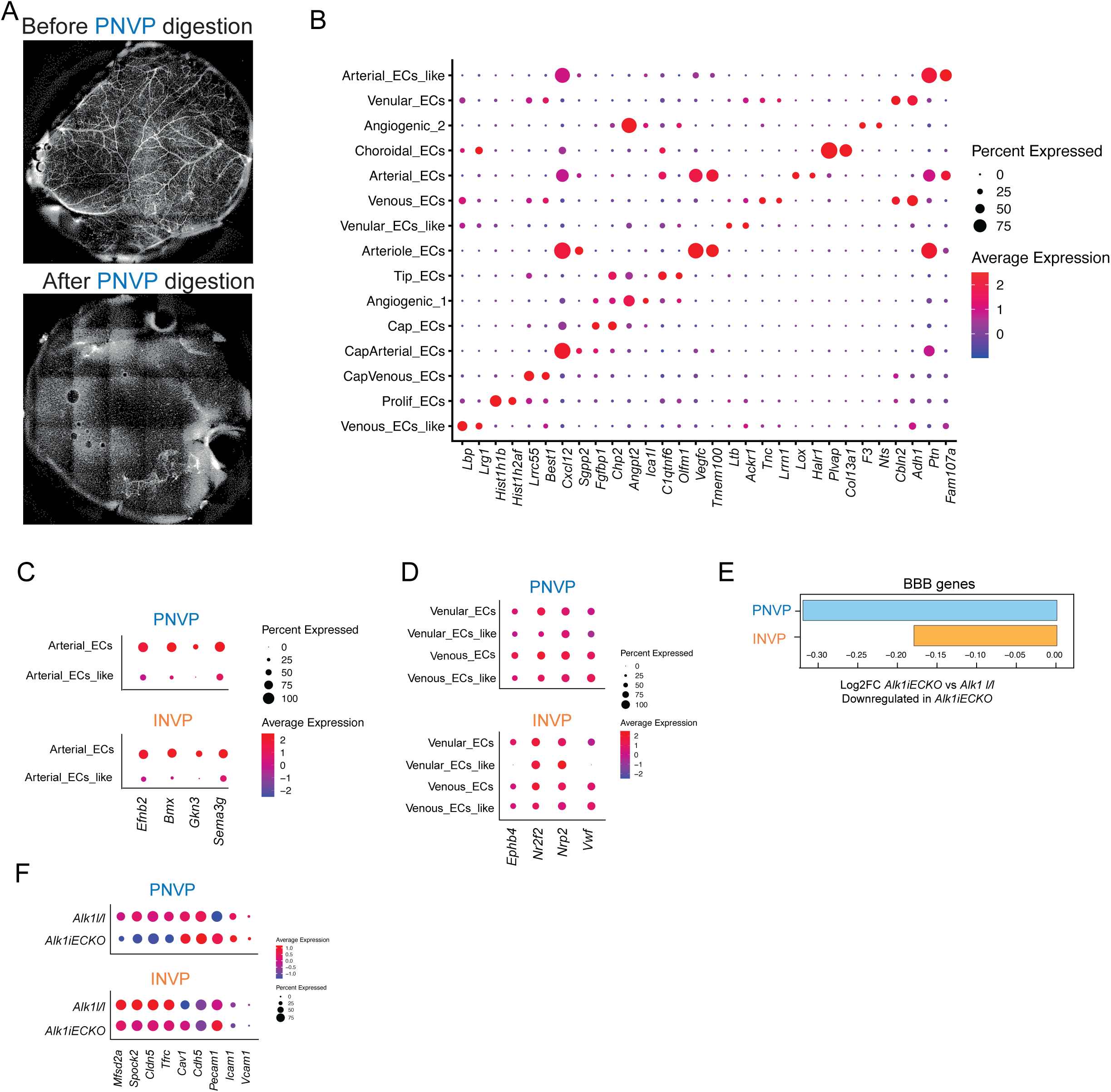
**A**, CD31 immunostaining nd wholemount imaging of P8 PNVP before (top) and after (bottom) PNVP digestion. **B**, Dot plot of top 2 markers expression of EC sub-clusters. **C-D**, Dot plot of selected arterial (C) and vein (D) EC markers. **E**, Barplot of VISION score log2FC of BBB gene expression in PNVP and INVP *Alk1iECKO* compared to control ECs. **F**, Dot plot of selected BBB EC markers.

**Supp. Figure 3.**
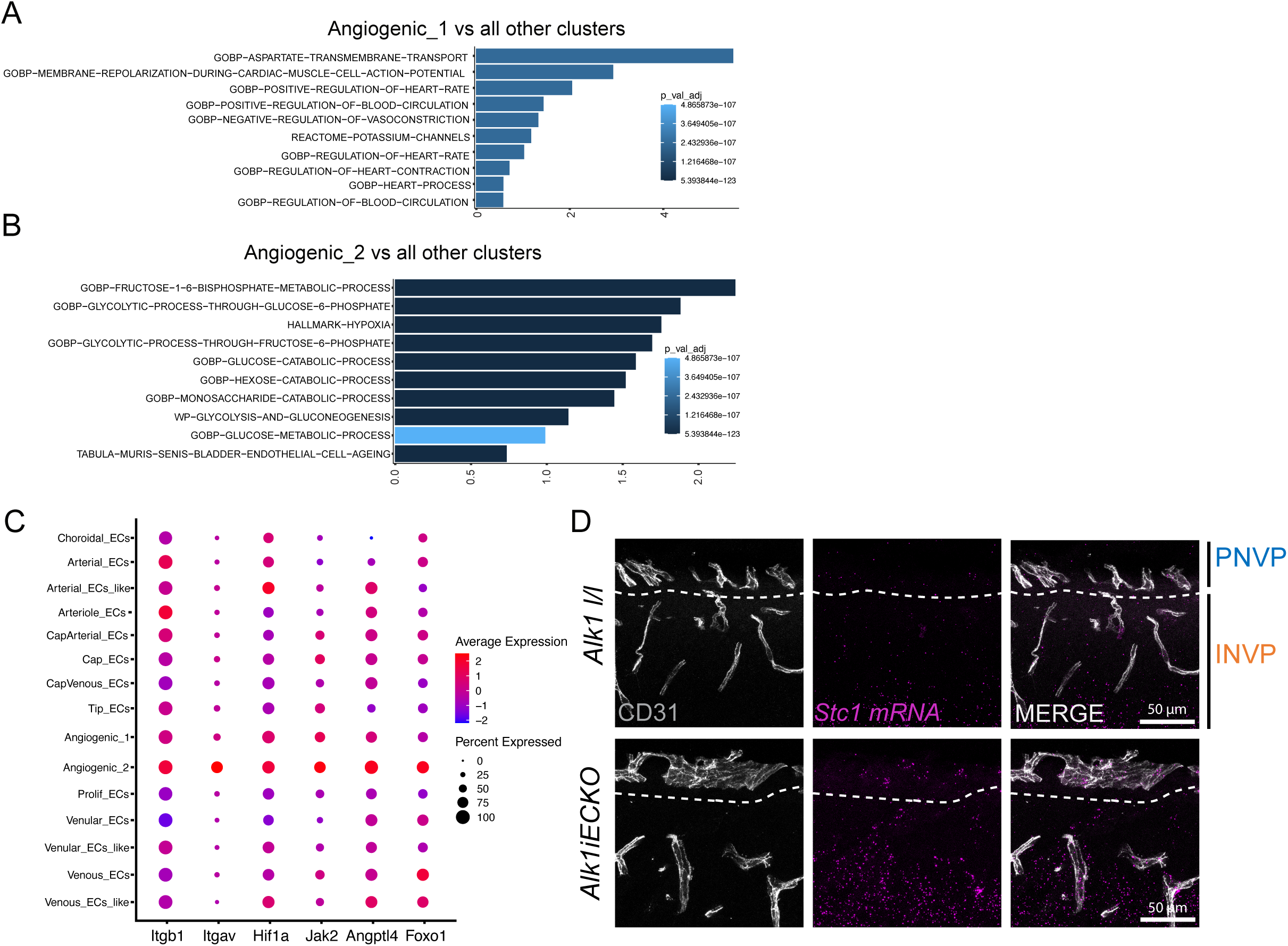
**A-B**, Barplot of VISION score log2FC of top 10 pathways upregulated in angiogenic 1 (A) and angiogenic 2 (B) ECs. **C**, Dot plot showing the expression of HHT-related genes across EC subclusters. **D**, *In situ* hybridization for *Stc1* mRNA using brain coronal section of P8 *Alk1iECKO* and control mice.

**Supp. Figure 4.**
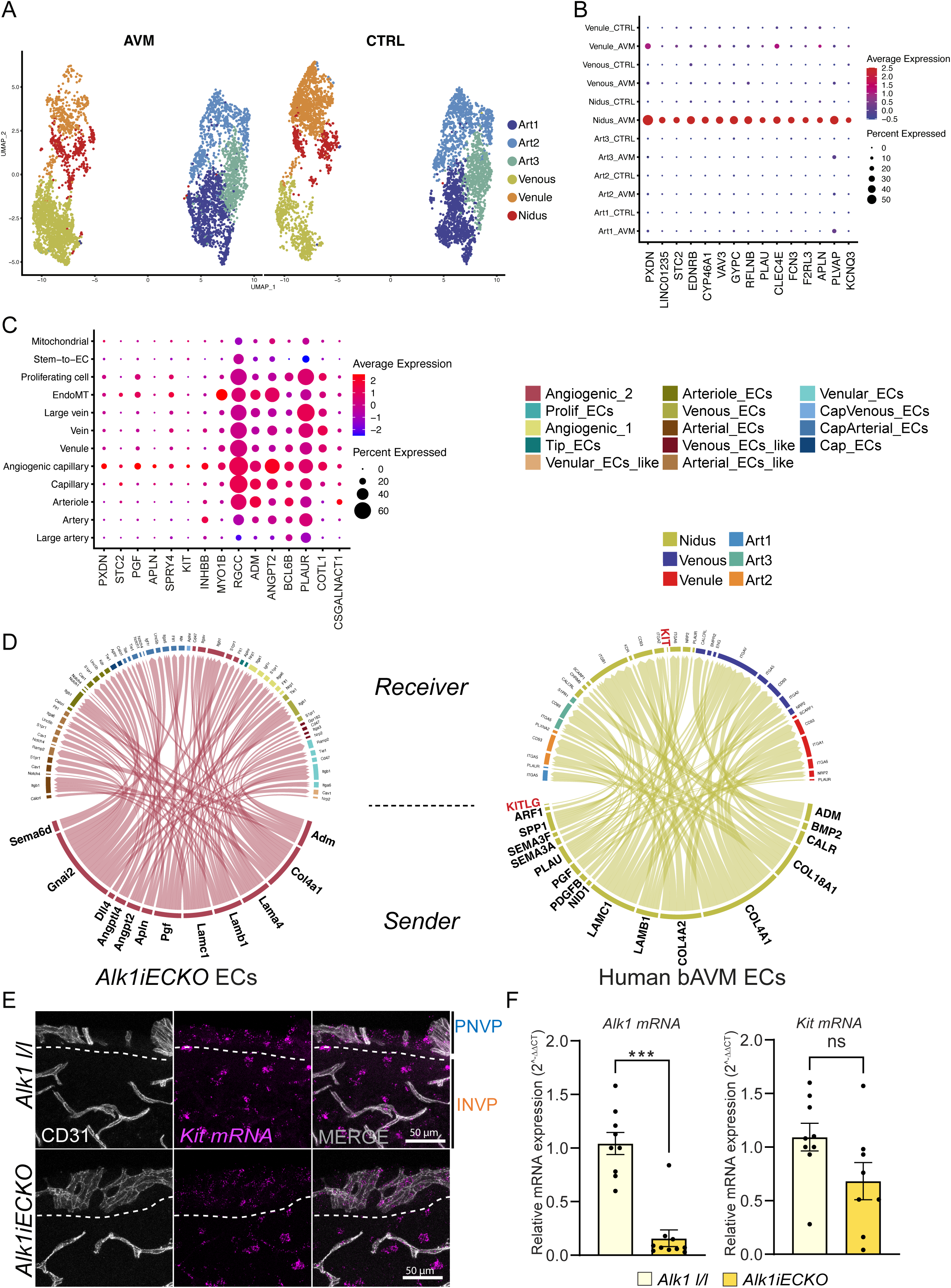
**A**, UMAP visualization of human brain AVM EC clusters (from Winkler E. et al 2022). **B**, Dot plot showing the expression of the top 10 markers in the human nidus EC subcluster (from Winkler E. et al 2022). **C**, Dot plot showing the expression of the top 10 markers in the human brain AVM angiogenic EC subcluster (from Walchli T. et al 2024). **D**, Circos plot representation of enriched ligand-receptor interactions in mouse (min.pct 0.5, min.z 0.5) and human (min.pct 0.2, min.z 0.5) EC clusters. **E**, *In situ* hybridization for *Kit* mRNA using brain coronal section of P8 *Alk1iECKO* and control mice. **F**, qPCR analysis of *Alk1* expression in isolated mouse lung ECs from *Alk1iECKO* and control mice. Each dot represents one mouse. Error bars represent means ± s.e.m, *\*\*\*P<0.001,* Mann-Whitney test or Unpaired t test (E) were performed.

**Supp. Figure 5.**
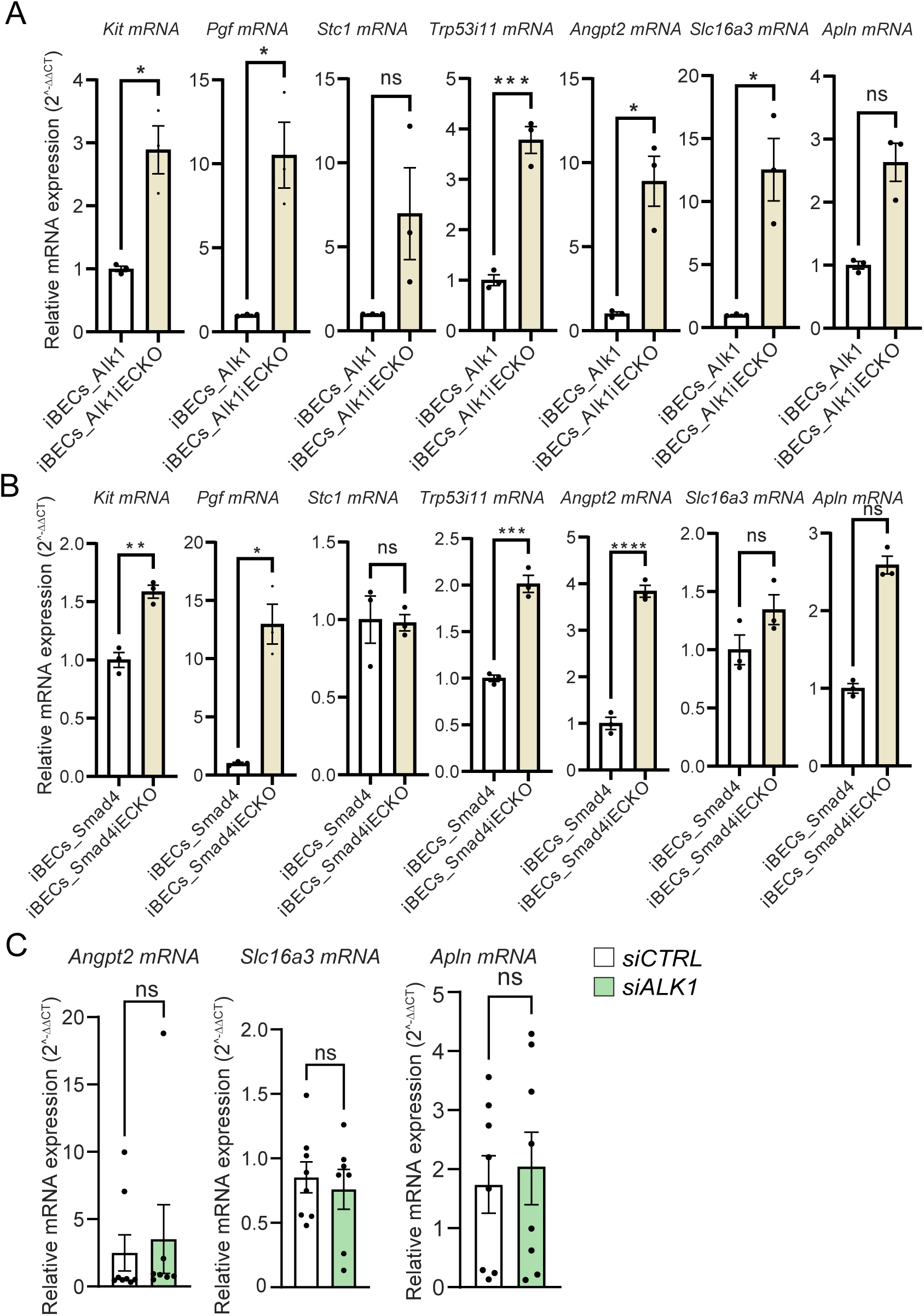
**A-B**,qPCR analysis of indicated gene expression in isolated mouse brain ECs from *Alk1iECKO* (A)*, Smad4iECKO* (B), and corresponding control mice. **C**, qPCR analysis of selected markers expression in HUVECs treated with ALK1 siRNA compared to control. Each dot represents one mouse (A, B, C) or experiment (D). Error bars represent means ± s.e.m, *\*P<0.05, **P<0.01, ***P<0.001, ****P<0.0001,* Mann-Whitney test, Welch’s t test or Unpaired t test (A, B C, D) were performed.

**Supp. Figure 6.**
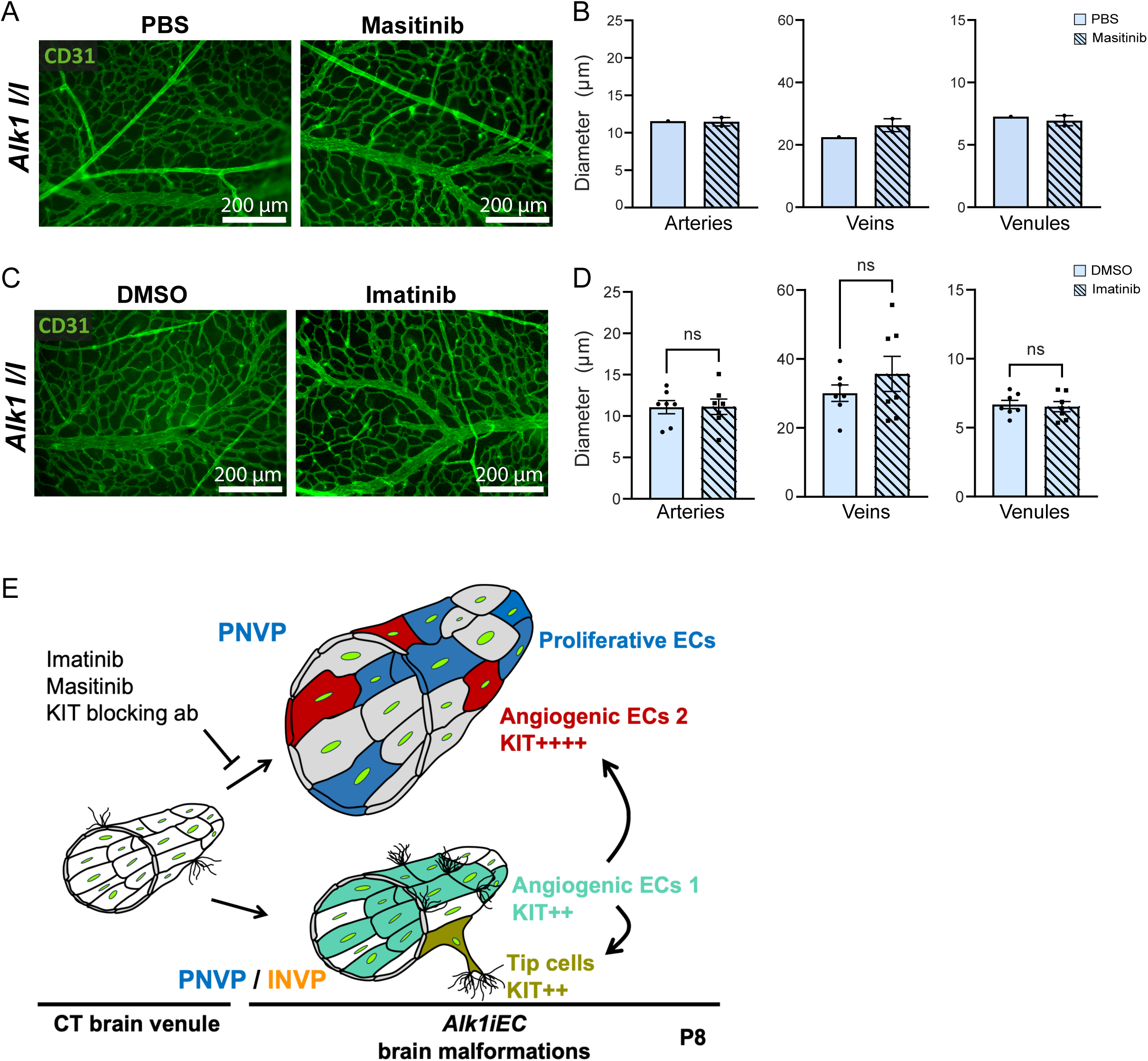
**A**, **C**, CD31 immunostaining of the PNVP of P8 *Alk1l/l* mice injected with imatinib (A), and masitinib (C), along with their corresponding vehicle controls. **B, D**, Quantification of vessel diameter in the PNVP of P8 *Alk1l/l* mice injected with the indicated inhibitor. Each dot represents one mouse. Error bars represent means ± s.e.m, Mann-Whitney test or Unpaired t test (B, D) were performed. **D**, Schematic model illustrating how 48 hours of Alk1 deletion induces the emergence of angiogenic 1 ECs in both PNVP and INVP. In the PNVP, angiogenic 1 ECs further differentiate into angiogenic 2 ECs, which drive AVM formation. Both angiogenic populations express KIT, with angiogenic 2 showing stronger expression. Pharmacological inhibition of KIT effectively prevents AVM development in this model. Each dot represents one mouse.

## References

1. Bideau A, Plauchu H, Brunet G, Robert J. Epidemiological investigation of Rendu-Osler disease in France: its geographical distribution and prevalence. Popul. 1989;44(1):3–22.

2. Kjeldsen AD, Vase P, Green A. Hereditary haemorrhagic telangiectasia: a population-based study of prevalence and mortality in Danish patients. Journal of Internal Medicine. 1999;245(1):31–39. doi:10.1046/j.1365-2796.1999.00398.x

3. McAllister KA, Grogg KM, Johnson DW, et al. Endoglin, a TGF-β binding protein of endothelial cells, is the gene for hereditary haemorrhagic telangiectasia type 1. Nat Genet. 1994;8(4):345–351. doi:10.1038/ng1294-345

4. Johnson DW, Berg JN, Baldwin MA, et al. Mutations in the activin receptor–like kinase 1 gene in hereditary haemorrhagic telangiectasia type 2. Nat Genet. 1996;13(2):189–195. doi:10.1038/ng0696-189

5. Prigoda NL, Savas S, Abdalla SA, et al. Hereditary haemorrhagic telangiectasia: mutation detection, test sensitivity and novel mutations. Journal of Medical Genetics. 2006;43(9):722–728. doi:10.1136/jmg.2006.042606

6. Howe JR, Roth S, Ringold JC, et al. Mutations in the SMAD4/DPC4 gene in juvenile polyposis. Science. 1998;280(5366):1086-1088. doi:10.1126/science.280.5366.1086

7. Wooderchak-Donahue WL, McDonald J, O’Fallon B, et al. BMP9 Mutations Cause a Vascular-Anomaly Syndrome with Phenotypic Overlap with Hereditary Hemorrhagic Telangiectasia. The American Journal of Human Genetics. 2013;93(3):530–537. doi:10.1016/j.ajhg.2013.07.004

8. Shovlin CL. Hereditary haemorrhagic telangiectasia: Pathophysiology, diagnosis and treatment. Blood Reviews. 2010;24(6):203–219. doi:10.1016/j.blre.2010.07.001

9. Brinjikji W, Iyer VN, Lanzino G, Thielen KR, Wood CP. Natural history of brain capillary vascular malformations in hereditary hemorrhagic telangiectasia patients. Journal of NeuroInterventional Surgery. 2017;9(1):26–28. doi:10.1136/neurintsurg-2015-012252

10. de Liyis BG, Arini AAIK, Karuniamaya CP, et al. Risk of intracranial hemorrhage in brain arteriovenous malformations: a systematic review and meta-analysis. J Neurol. 2024;271(5):2274–2284. doi:10.1007/s00415-024-12235-1

11. Josephson CB, Rosenow F, Salman RAS. Intracranial Vascular Malformations and Epilepsy. Seminars in Neurology. 2015;35:223-234. doi:10.1055/s-0035-1552621

12. Tabosh TA, Tarrass MA, Tourvieilhe L, Guilhem A, Dupuis-Girod S, Bailly S. Hereditary hemorrhagic telangiectasia: from signaling insights to therapeutic advances. J Clin Invest. 2024;134(4). doi:10.1172/JCI176379

13. Sugiyama T, Grasso G, Torregrossa F, Fujimura M. Current Concepts and Perspectives on Brain Arteriovenous Malformations: A Review of Pathogenesis and Multidisciplinary Treatment. World Neurosurgery. 2022;159:314–326. doi:10.1016/j.wneu.2021.07.106

14. Tual-Chalot S, Mahmoud M, Allinson KR, et al. Endothelial Depletion of Acvrl1 in Mice Leads to Arteriovenous Malformations Associated with Reduced Endoglin Expression. PLOS ONE. 2014;9(6):12.

15. Mahmoud M, Allinson KR, Zhai Z, et al. Pathogenesis of Arteriovenous Malformations in the Absence of Endoglin. Circulation Research. 2010;106(8):1425–1433. doi:10.1161/CIRCRESAHA.109.211037

16. Ola R, Künzel SH, Zhang F, et al. SMAD4 Prevents Flow Induced Arteriovenous Malformations by Inhibiting Casein Kinase 2. Circulation. 2018;138(21):2379–2394. doi:10.1161/CIRCULATIONAHA.118.033842

17. Benn A, Alonso F, Mangelschots J, Génot E, Lox M, Zwijsen A. BMP-SMAD1/5 Signaling Regulates Retinal Vascular Development. Biomolecules. 2020;10(3):488. doi:10.3390/biom10030488

18. Park H, Furtado J, Poulet M, et al. Defective Flow-Migration Coupling Causes Arteriovenous Malformations in Hereditary Hemorrhagic Telangiectasia. Circulation. 2021;144(10):805–822. doi:10.1161/CIRCULATIONAHA.120.053047

19. Lee HW, Xu Y, He L, et al. Role of Venous Endothelial Cells in Developmental and Pathologic Angiogenesis. Circulation. 2021;144(16):1308–1322. doi:10.1161/CIRCULATIONAHA.121.054071

20. Drapé E, Anquetil T, Larrivée B, Dubrac A. Brain arteriovenous malformation in hereditary hemorrhagic telangiectasia: Recent advances in cellular and molecular mechanisms. Frontiers in Human Neuroscience. 2022;16. Accessed December 1, 2022. https://www.frontiersin.org/articles/10.3389/fnhum.2022.1006115

21. Ola R, Dubrac A, Han J, et al. PI3 kinase inhibition improves vascular malformations in mouse models of hereditary haemorrhagic telangiectasia. Nat Commun. 2016;7:13650. doi:10.1038/ncomms13650

22. Park S, DiMaio TA, Liu W, Wang S, Sorenson CM, Sheibani N. Endoglin regulates the activation and quiescence of endothelium by participating in canonical and non-canonical TGF-β signaling pathways. Journal of Cell Science. 2013;126(6):1392–1405. doi:10.1242/jcs.117275

23. Marziano C, Genet G, Hirschi KK. Vascular endothelial cell specification in health and disease. Angiogenesis. 2021;24(2):213–236. doi:10.1007/s10456-021-09785-7

24. Winkler EA, Kim CN, Ross JM, et al. A single-cell atlas of the normal and malformed human brain vasculature. Science. 2022;375(6584):eabi7377. doi:10.1126/science.abi7377

25. Wälchli T, Ghobrial M, Schwab M, et al. Single-cell atlas of the human brain vasculature across development, adulthood and disease. Nature. 2024;632(8025):603-613. doi:10.1038/s41586-024-07493-y

26. Genet G, Genet N, Paila U, et al. Induced Endothelial Cell Cycle Arrest Prevents Arteriovenous Malformations in Hereditary Hemorrhagic Telangiectasia. Circulation. 2024;149(12):944–962. doi:10.1161/CIRCULATIONAHA.122.062952

27. Yang Y, Wu X, Zhao Y, et al. Arterial-Lymphatic-Like Endothelial Cells Appear in Hereditary Hemorrhagic Telangiectasia 2 and Contribute to Vascular Leakage and Arteriovenous Malformations. Circulation. 2025;151(5):299–317. doi:10.1161/CIRCULATIONAHA.124.070925

28. Hao Y, Stuart T, Kowalski MH, et al. Dictionary learning for integrative, multimodal and scalable single-cell analysis. Nat Biotechnol. 2024;42(2):293–304. doi:10.1038/s41587-023-01767-y

29. Coelho-Santos V, Shih AY. Postnatal development of cerebrovascular structure and the neurogliovascular unit. Wiley Interdiscip Rev Dev Biol. 2020;9(2):e363. doi:10.1002/wdev.363

30. Vanlandewijck M, He L, Mäe MA, et al. A molecular atlas of cell types and zonation in the brain vasculature. Nature. 2018;554(7693):475–480. doi:10.1038/nature25739

31. Lendahl U, Nilsson P, Betsholtz C. Emerging links between cerebrovascular and neurodegenerative diseases—a special role for pericytes. EMBO reports. 2019;20(11):e48070. doi:10.15252/embr.201948070

32. Larrivée B, Prahst C, Gordon E, et al. ALK1 Signaling Inhibits Angiogenesis by Cooperating with the Notch Pathway. Developmental Cell. 2012;22(3):489–500. doi:10.1016/j.devcel.2012.02.005

33. Winkler EA, Birk H, Burkhardt JK, et al. Reductions in brain pericytes are associated with arteriovenous malformation vascular instability. J Neurosurg. 2018;129(6):1464-1474. doi:10.3171/2017.6.JNS17860

34. Brown RD, Wiebers DO, Torner JC, O’Fallon WM. Frequency of intracranial hemorrhage as a presenting symptom and subtype analysis: a population-based study of intracranial vascular malformations in Olmsted County, Minnesota. Journal of Neurosurgery. 1996;85(1):29–32. doi:10.3171/jns.1996.85.1.0029

35. Ricciardelli AR, Robledo A, Fish JE, Kan PT, Harris TH, Wythe JD. The Role and Therapeutic Implications of Inflammation in the Pathogenesis of Brain Arteriovenous Malformations. Biomedicines. 2023;11(11):2876. doi:10.3390/biomedicines11112876

36. Tual-Chalot S, Oh P, Arthur H. Mouse Models of Hereditary Haemorrhagic Telangiectasia: Recent Advances and Future Challenges. Frontiers in Genetics. 2015;6. Accessed May 30, 2022. https://www.frontiersin.org/article/10.3389/fgene.2015.00025

37. Cerdà P, Castillo SD, Aguilera C, et al. New genetic drivers in hemorrhagic hereditary telangiectasia. Eur J Intern Med. 2024;119:99–108. doi:10.1016/j.ejim.2023.08.024

38. Raredon MSB, Yang J, Garritano J, et al. Computation and visualization of cell–cell signaling topologies in single-cell systems data using Connectome. Sci Rep. 2022;12(1):4187. doi:10.1038/s41598-022-07959-x

39. Broudy VC. Stem Cell Factor and Hematopoiesis. Blood. 1997;90(4):1345-1364. doi:10.1182/blood.V90.4.1345

40. Sheikh E, Tran T, Vranic S, Levy A, Bonfil RD. Role and significance of c-KIT receptor tyrosine kinase in cancer: A review. Bosn J Basic Med Sci. 2022;22(5):683–698. doi:10.17305/bjbms.2021.7399

41. Klug LR, Corless CL, Heinrich MC. Inhibition of KIT Tyrosine Kinase Activity: Two Decades After the First Approval. J Clin Oncol. 2021;39(15):1674–1686. doi:10.1200/JCO.20.03245

42. Zhou X, Pucel JC, Nomura-Kitabayashi A, et al. ANG2 Blockade Diminishes Proangiogenic Cerebrovascular Defects Associated With Models of Hereditary Hemorrhagic Telangiectasia. *Arteriosclerosis*, Thrombosis, and Vascular Biology. 2023;43(8):1384–1403. doi:10.1161/ATVBAHA.123.319385

43. Dubreuil P, Letard S, Ciufolini M, et al. Masitinib (AB1010), a Potent and Selective Tyrosine Kinase Inhibitor Targeting KIT. PLoS One. 2009;4(9):e7258. doi:10.1371/journal.pone.0007258

44. Allard B, Jacoberger-Foissac C, Cousineau I, et al. Adenosine A2A receptor is a tumor suppressor of NASH-associated hepatocellular carcinoma. Cell Rep Med. 2023;4(9):101188. doi:10.1016/j.xcrm.2023.101188

45. Baeyens N, Larrivée B, Ola R, et al. Defective fluid shear stress mechanotransduction mediates hereditary hemorrhagic telangiectasia. Journal of Cell Biology. 2016;214(7):807–816. doi:10.1083/jcb.201603106

46. Komiyama M, Terada A, Ishiguro T, et al. Neuroradiological Manifestations of Hereditary Hemorrhagic Telangiectasia in 139 Japanese Patients. Neurologia medico-chirurgica. 2015;55(6):479–486. doi:10.2176/nmc.oa.2015-0040

47. Kim KL, Meng Y, Kim JY, Baek EJ, Suh W. Direct and differential effects of stem cell factor on the neovascularization activity of endothelial progenitor cells. Cardiovascular Research. 2011;92(1):132–140. doi:10.1093/cvr/cvr161

48. Shan H jian, Jiang K, Zhao M zhi, et al. SCF/c-Kit-activated signaling and angiogenesis require Gαi1 and Gαi3. Int J Biol Sci. 2023;19(6):1910–1924. doi:10.7150/ijbs.82855

49. Feng T, Gao Z, Kou S, et al. No Evidence for Erythro-Myeloid Progenitor-Derived Vascular Endothelial Cells in Multiple Organs. Circulation Research. 2020;127(10):1221–1232. doi:10.1161/CIRCRESAHA.120.317442

50. Lennartsson J, Rönnstrand L. Stem Cell Factor Receptor/c-Kit: From Basic Science to Clinical Implications. Physiological Reviews. 2012;92(4):1619–1649. doi:10.1152/physrev.00046.2011

51. Pedersen M, Löfstedt T, Sun J, Holmquist-Mengelbier L, Påhlman S, Rönnstrand L. Stem cell factor induces HIF-1alpha at normoxia in hematopoietic cells. Biochem Biophys Res Commun. 2008;377(1):98–103. doi:10.1016/j.bbrc.2008.09.102

52. Kierans SJ, Taylor CT. Regulation of glycolysis by the hypoxia-inducible factor (HIF): implications for cellular physiology. The Journal of Physiology. 2021;599(1):23–37. doi:10.1113/JP280572

53. Szwed A, Kim E, Jacinto E. Regulation and metabolic functions of mTORC1 and mTORC2. Physiological Reviews. 2021;101(3):1371–1426. doi:10.1152/physrev.00026.2020

